# Efficient and precise single-cell reference atlas mapping with Symphony

**DOI:** 10.1101/2020.11.18.389189

**Authors:** Joyce B. Kang, Aparna Nathan, Fan Zhang, Nghia Millard, Laurie Rumker, D. Branch Moody, Ilya Korsunsky, Soumya Raychaudhuri

**Affiliations:** Center for Data Sciences, Brigham and Women’s Hospital, Boston, MA, USA; Division of Genetics, Department of Medicine, Brigham and Women’s Hospital and Harvard Medical School, Boston, MA, USA; Division of Rheumatology, Inflammation, and Immunity, Department of Medicine, Brigham and Women’s Hospital and Harvard Medical School, Boston, MA, USA; Department of Biomedical Informatics, Harvard Medical School, Boston, MA, USA; Program in Medical and Population Genetics, Broad Institute of MIT and Harvard, Cambridge, MA, USA; Versus Arthritis Centre for Genetics and Genomics, Centre for Musculoskeletal Research, Manchester Academic Health Science Centre, The University of Manchester, Manchester, UK

**Keywords:** single-cell genomics, scRNA-seq, reference mapping, annotation

## Abstract

Recent advances in single-cell technologies and integration algorithms make it possible to construct comprehensive reference atlases encompassing many donors, studies, disease states, and sequencing platforms. Much like mapping sequencing reads to a reference genome, it is essential to be able to map query cells onto complex, multimillion-cell reference atlases to rapidly identify relevant cell states and phenotypes. We present Symphony (https://github.com/immunogenomics/symphony), an algorithm for building integrated reference atlases of millions of cells in a convenient, portable format that enables efficient query mapping within seconds. Symphony localizes query cells within a stable low-dimensional reference embedding, facilitating reproducible downstream transfer of reference-defined annotations to the query. We demonstrate the power of Symphony by (1) mapping a multi-donor, multi-species query to predict pancreatic cell types, (2) localizing query cells along a developmental trajectory of human fetal liver hematopoiesis, and (3) inferring surface protein expression with a multimodal CITE-seq atlas of memory T cells.

## Introduction

Advancements in single-cell RNA-sequencing (scRNA-seq) have launched an era in which individual studies can routinely profile 10^4^-10^6^ cells^1–3^, and multimillion-cell datasets are already emerging^4,5^.Single-cell resolution enables the discovery and refinement of cell states across diverse clinical and biological contexts^6–11^. To date, most studies redefine cell states from scratch, making it difficult to compare results across studies and thus hampering reproducibility. Coordinated large-scale efforts, exemplified by the Human Cell Atlas (HCA)^12^, aim to establish comprehensive and well-annotated reference datasets comprising millions of cells that capture the broad spectrum of cell states. Building these reference atlases requires integrating multiple datasets that may have been collected under different technical and biological conditions. Hence, reference construction requires application of one of many recently developed single-cell integration algorithms^13–19^. Our group previously developed Harmony^15^, a fast, accurate, and well-reviewed method^20^ that is able to explicitly model complex study design, a property that makes it suitable for integrating complex datasets into reference atlases^21–24^. The potential to define common cell states using reference maps has already been demonstrated^25,26^. For example, we built an integrated reference of ∼80,000 single-cell profiles of fibroblasts from human lung, synovium, salivary gland, and intestine and successfully mapped fibroblasts from human skin and mouse synovium, lung, and intestine to analyze conserved states across tissues and species^25^. Once such reference atlases are painstakingly constructed, interpretation of new datasets requires the ability to quickly map single-cell profiles into these reference atlases. This enables interpretation of new datasets by transferring annotations and metadata of interest from nearby reference cells.

Fast mapping of query cells against a large, stable reference is a well-recognized open problem^27^ and active area of research^28–30^. One inefficient but accurate approach to project reference and query cells into a joint embedding is to integrate both sets of cells together *de novo*, resulting in what might be considered a “gold standard” embedding. While this approach is reasonable for relatively small reference datasets, it is intractable for atlas-sized references with millions of cells. It requires users to “rebuild” the reference for each analysis, which may be computationally challenging and require administratively cumbersome exchanges of large-scale datasets. Furthermore, *de novo* integration may corrupt the reference embedding once a reference is carefully constructed and annotated. It is instead preferable to freeze the reference when mapping new query cells onto it.

Here, we define reference mapping to mean placing query cells within the same embedding as integrated reference cells without requiring access to the raw data on all individual reference cells.Importantly, this embedding does not take advantage of any particular annotation, such as cell type labels, which may be refined or updated over time. This is in contrast to automated cell type classifiers, such as scmap^31^, which assign rigid annotations based on reference datasets in a supervised manner. Reference mapping approaches introduced so far include Seurat v4^30^, which is compatible with Seurat integration^18^, and scArches, which is compatible with autoencoders such as scANVI^32^ and trVAE^33^.These approaches separate reference building, which integrates datasets in the reference into a low-dimensional embedding, from query mapping, which uses a compressed version of the reference to efficiently map cells into the reference embedding. They further contrast with *de novo* integration methods like BBKNN^34^, Seurat v3^18^, and Harmony^17^, which enable reference building but are slow and require access to the raw data and batch information on individual reference cells. High-quality reference mapping requires both a framework to efficiently store an integrated reference, and a fast and accurate procedure to map query datasets.

An ideal reference mapping algorithm must meet four key requirements. First, similar to *de novo* integration algorithms, they must be able to remove confounding signals due to complex study design in both the reference and query. In addition, they must be able to scale to large datasets, map with high accuracy, and enable inference of diverse query cell annotations based on reference cells. We present Symphony, a novel algorithm to compress a large, integrated reference and map query cells to a precise location in the reference embedding within seconds. Through multiple real-world dataset analyses, we show that Symphony can enable accurate downstream inference of cell type, developmental trajectory position, and protein expression, even when the query itself contains complex confounding technical and biological effects.

## Results

### Symphony compresses an integrated reference for efficient query mapping

Symphony comprises two main algorithms: reference compression and mapping (**Methods, Fig. S1a**). Symphony *reference compression* captures and structures information from multiple reference datasets into an integrated and concise format that can subsequently be used to map query cells (**Fig. 1a-b**).Symphony builds upon the linear mixture model framework first introduced by Harmony^17^. Briefly, in a low-dimensional embedding, such as principal component analysis (PCA), the model represents cell states as soft clusters, in which a cell’s identity is defined by probabilistic assignments across one or more clusters. For *de novo* integration of the reference, cells are iteratively assigned soft cluster memberships, which are used as weights in a linear mixture model to remove unwanted covariate-dependent effects. To store the reference efficiently without saving information on individual reference cells, Symphony computes summary statistics learned in the low-dimensional space (**Fig. 1b, Methods**), returning computationally efficient data structures containing the “minimal reference elements” needed to map new cells. These include the means and standard deviations used to scale the genes, the gene loadings from PCA (or another low dimensional projection, e.g. canonical correlation analysis [CCA]) on the reference cells, soft-cluster centroids from the integrated reference, and two “compression terms” (a *k* x 1 vector and *k* x *d* matrix, where *k* is the number of clusters and *d* is the dimensionality of the embedding) (**Methods, Supplementary Equations, Fig. S1b**).

**Figure 1.**
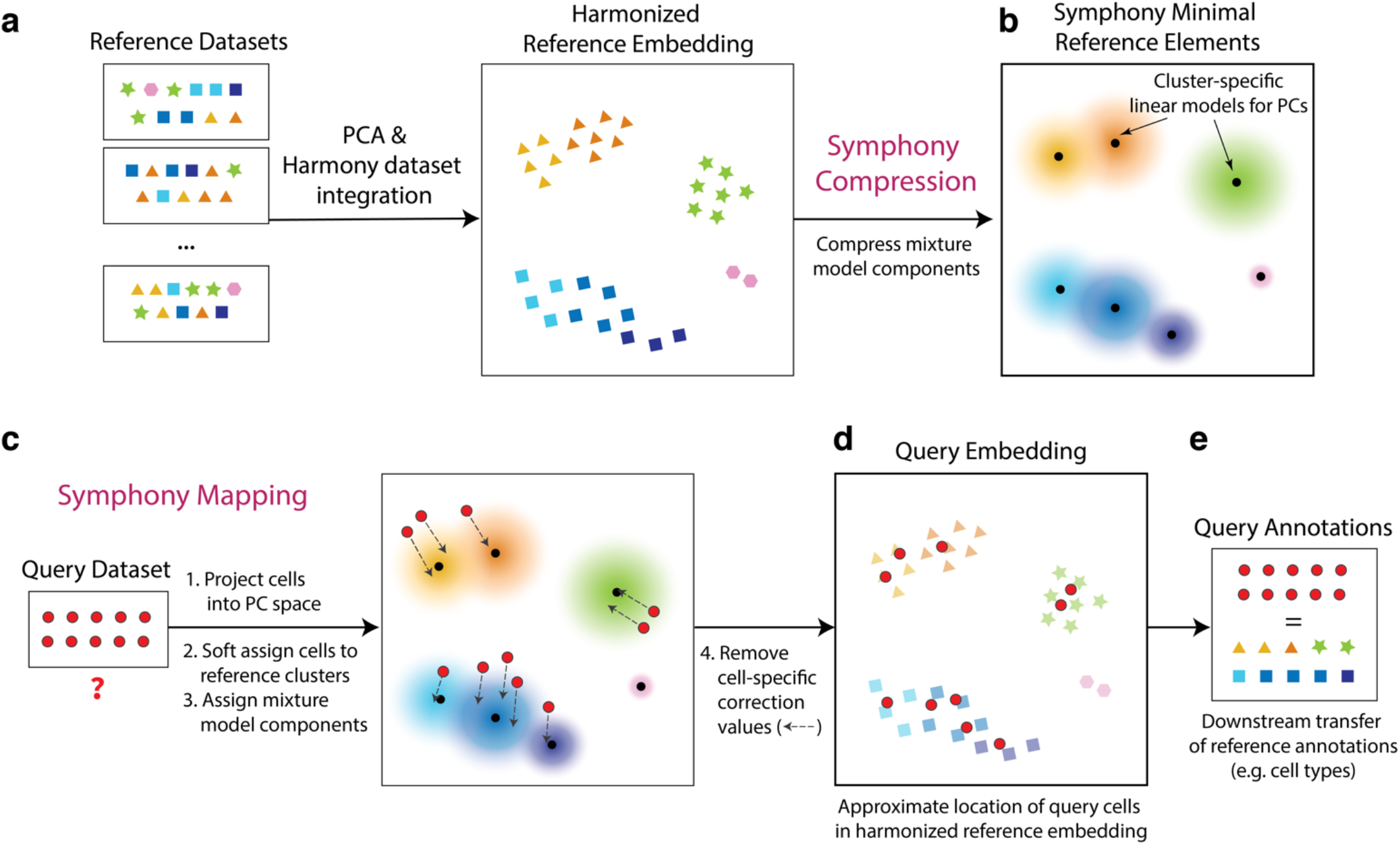
Symphony Overview. Symphony comprises two algorithms: Symphony compression **(a-b)** and Symphony mapping **(c-d). (a)** To construct a reference atlas, cells from multiple datasets are embedded in a lower-dimensional space (e.g. PCA), in which dataset integration (Harmony) is performed to remove dataset-specific effects. Shape indicates distinct cell types, and color indicates finer-grained cell states. **(b)** Symphony compression represents the information captured within the harmonized reference in a concise, portable format based on computing summary statistics for the reference-dependent components of the linear mixture model. Symphony returns the minimal reference elements needed to efficiently map new query cells to the reference. **(c)** Given an unseen query dataset and compressed reference, Symphony mapping precisely localizes the query cells to their appropriate locations within the integrated reference embedding **(d)**. Reference cell locations do not change during mapping. **(e)** The resulting joint embedding can be used for downstream transfer of reference-defined annotations to the query cells. See Fig. S1.

To map new query cells to the compressed reference, we apply Symphony *mapping*. The algorithm approximates integration of reference and query cells *de novo* (**Methods**), but uses only the minimal reference elements to compute the mapping (**Fig. S1c**). First, Symphony projects query gene expression profiles into the same uncorrected low-dimensional space as the reference cells (e.g. PCs), using the saved scaling parameters and reference gene loadings (**Fig. 1c**). Second, Symphony computes soft cluster assignments for the query cells based on proximity to the reference cluster centroids. Finally, to correct unwanted user-specified technical and biological effects in the query data, Symphony assumes the soft cluster assignments from the previous step and uses stored mixture model components to estimate and regress out the query batch effects (**Fig. 1d**). Importantly, the reference cell embedding remains stable during mapping. Embedding the query within the reference coordinates enables downstream transfer of annotations from reference cells to query cells, including discrete cell type classifications, quantitative cell states (e.g. position along a trajectory), or expression of missing genes or proteins (**Fig. 1e**).

### Symphony approximates *de novo* integration of PBMCs without reintegration of reference datasets

As we demonstrate in the **Methods**, Symphony is equivalent to running *de novo* Harmony integration if three conditions are met: (I) all cell states represented in the query data set are captured by the reference dataset, (II) the number of query cells is much smaller than the number of reference cells, and (III) the query dataset has a design matrix that is independent of reference datasets (i.e. non-overlapping batches in reference and query). As the scope of available single-cell atlases continues to grow, it is reasonable to assume that reference datasets are large and all-inclusive, making conditions (I) and (II) well-supported. Condition (III) is also typically met if the query data was generated in separate experiments from the reference.

To demonstrate that Symphony mapping closely approximates running *de novo* integration on all cells, we applied Symphony to 20,792 peripheral blood mononuclear cells (PBMCs) assayed with three different 10x technologies: 3’v1, 3’v2, and 5’. We performed three mapping experiments. For each, we built an integrated Symphony reference from two technologies, then mapped the third technology as a query. The resulting Symphony embeddings were compared to a gold standard embedding obtained by running Harmony on all three datasets together. Visually, we found that the Symphony embedding for each mapping experiment (**Fig. 2a**) closely reproduced the overall structure and cell type information of the gold standard embedding (**Fig. 2b**). To quantitatively assess the degrees of dataset mixing we use the Local Inverse Simpson’s Index (LISI)^17^ metric. For a given categorical label assigned to each cell (in this case, technology), LISI indicates the effective number of categories represented in the local neighborhood of each cell; higher LISI scores correspond to better mixing of cells across batches. LISI scores in Symphony embeddings (mean LISI 2.16, 95% CI [2.16, 2.17]) and *de novo* integration embeddings (mean LISI 2.14, 95% CI [2.13, 2.15]) were nearly identical (**Fig. 2c, Methods**).

**Figure 2.**
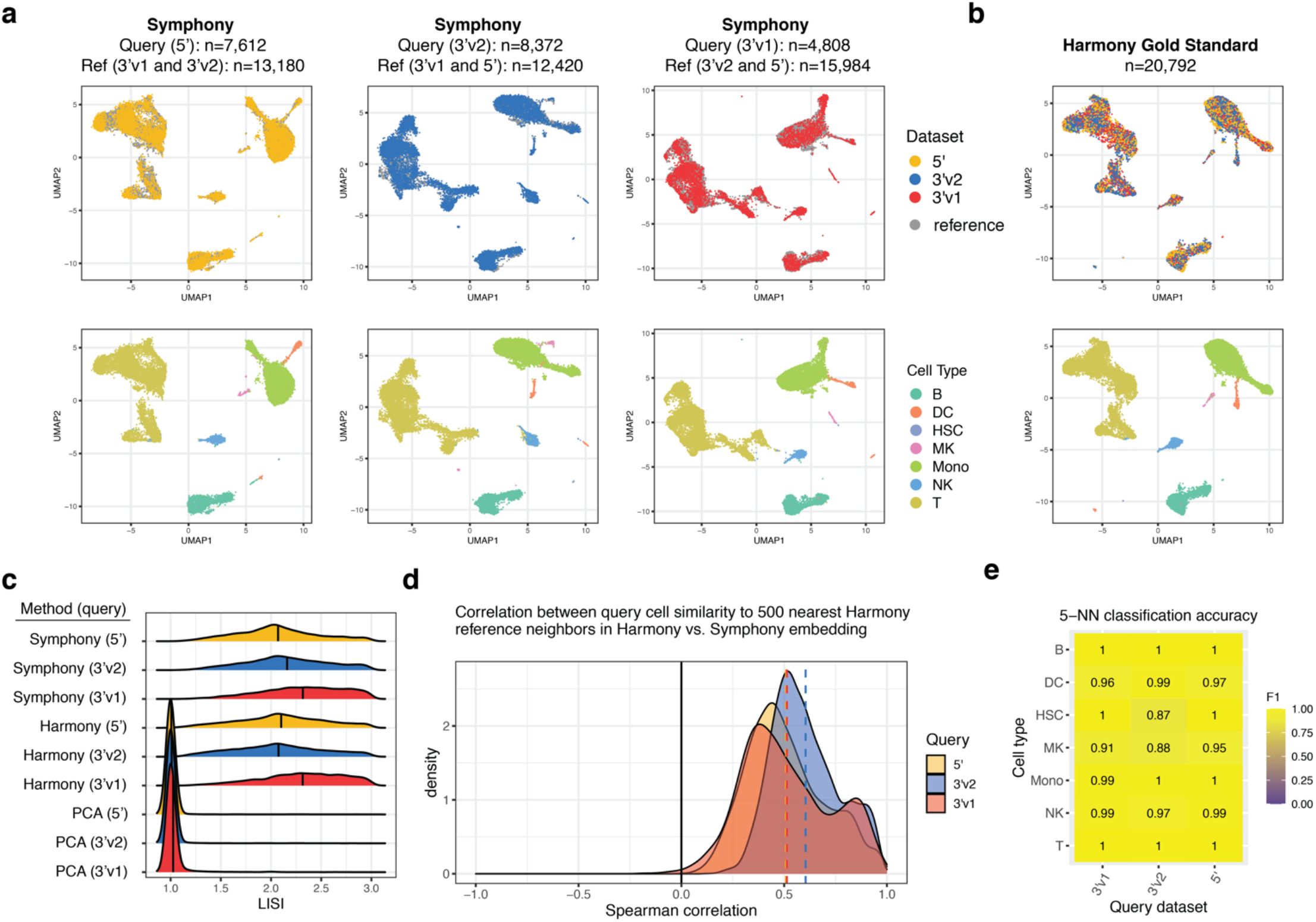
Symphony approximates *de novo* integration without reintegration of the reference cells. Three PBMC datasets were sequenced with different 10x protocols: 5’ (yellow, n=7,697 cells), 3’v2 (blue, n=8,380 cells), and 3’v1 (red, n=4,809 cells). We ran Symphony three times, each time mapping one dataset onto a reference built from integrating the other two. **(a)** Symphony embeddings generated across the three mapping experiments (columns). Top row: cells colored by query (yellow, blue, or red) or reference (gray), with query cells plotted in front. Bottom row: cells colored by cell type: B cell (B), dendritic cell (DC), hematopoietic stem cell (HSC), megakaryocyte (MK), monocyte (Mono), natural killer cell (NK), or T cell (T), with query cells plotted in front. **(b)** For comparison, gold standard *de novo* Harmony embedding colored by dataset (top) and cell type (bottom). **(c)** Distribution of technology LISI scores for query cell neighborhoods in the Symphony, gold standard, and a standard PCA embeddings on all cells. **(d)** Distribution of k-NN-corr (Spearman correlation between the similarities between the neighbor-pairs in the Harmony embedding and the similarities between the same neighbor-pairs in the Symphony embedding) across query cells for k=500, colored by query dataset. **(e)** Classification accuracy as measured by cell type F1 scores for query cell type annotation using 5-NN on the Symphony embedding. See Fig. S2.

To directly assess similarity of the local neighborhood structures, we computed the correlation between the local neighborhood adjacency graphs generated by Symphony and *de novo* integration. We define a new metric called k-nearest-neighbor correlation (k-NN-corr), which quantifies how well the local neighborhood structure in a given embedding is preserved in an alternative embedding by looking at the correlation of neighbor cells sorted by distance (**Fig. S2a-e**). Anchoring on each query cell, we calculate (1) the pairwise similarities to its *k* nearest reference neighbors in the gold standard embedding and (2) the similarities between the same query-reference neighbor pairs in the alternate embedding (**Methods**), then calculate the Spearman correlation between (1) and (2). k-NN-corr ranges from -1 to +1, where +1 indicates a perfectly preserved sorted ordering of neighbors. We find that for *k*=500, the Symphony embeddings produce a k-NN-corr >0.4 for 77.3% of cells (and positive k-NN-corr for 99.9% of cells), demonstrating that Symphony not only maps query cells to the correct broad cluster but also preserves the distance relationships between nearby cells in the same local region (**Fig. 2d)**.As a comparison, we calculated k-NN-corr for a simple PC projection of the query cells (with no correction step) using the original reference gene loadings prior to integration and observed significantly lower correlations (Wilcoxon signed-rank p<2.2e-16), with k-NN-corr >0.4 for 39.9% of cells (**Fig. S2f**).

### Symphony enables accurate cell type classification of PBMCs across technologies

If Symphony is effective, then cells should be mapped close to cells of the same cell type, enabling accurate cell type classification. To test this, we performed post-mapping query cell type classification in the 10x PBMCs example from above. We used a 5-NN classifier to annotate query cells across 7 cell types based on the nearest reference cells in the harmonized embedding and compared the predictions to the ground truth labels assigned *a priori* with lineage-specific marker genes (**Methods, Table S2**). Across all three experiments, predictions using the Symphony embeddings achieved 99.5% accuracy overall, with a median cell type F1-score (harmonic mean of precision and recall, ranging from 0 to 1) of 0.99 (**Fig. 2e, Table S3**). This indicates that Symphony appropriately localizes query cells in harmonized space to enable the accurate transfer of cell type labels.

Automatic cell type classification represents an open area of research^31,35–38^. Existing supervised classifiers assign a limited set of labels to new cells based on training data and/or marker genes. To benchmark Symphony-powered downstream inference against existing classifiers, we followed the same procedure as a benchmarking analysis in Abdelaal et al. (2019)^35^. The benchmark compared 22 cell type classifiers on the PbmcBench dataset consisting of two PBMC samples sequenced using 7 different protocols^39^. For each protocol train-test pair (42 experiments) and donor train-test pair (additional 6 experiments) (**Methods**), we built a Symphony reference from the training dataset then mapped the test dataset. We used the resulting harmonized feature embedding to predict query cell types using three downstream models: 5-NN, SVM with radial kernel, and multinomial logistic regression. The Symphony-based classifiers achieve consistently high cell type F1-scores (average median F1 of 0.79-0.83) comparable to the top three supervised classifiers for this benchmark (scmapcell, singleCellNet, and SCINA, average median F1 of 0.77-0.83) (**Fig. S3a**). Notably, as discussed in Abdelaal et al., the median F1-score alone can be misleading given that some classifiers (including SCINA) leave low-confidence cells as “unclassified”, whereas we used Symphony to assign a label to every cell. This benchmark is also arguably suboptimal in that the reference in each experiment is comprised of a single dataset (no reference integration involved).

### Symphony maps against a large reference within seconds

To demonstrate scalability to large reference atlases, we evaluated Symphony’s computational speed. We downsampled a large memory T cell dataset^40^ to create benchmark reference datasets with 20,000, 50,000, 100,000, 250,000, and 500,000 cells (from 12, 30, 58, 156, and 259 donors, respectively). Against each reference, we mapped three different-sized queries: 1,000, 10,000, and 100,000 cells (from 1, 6, and 64 donors) and measured total elapsed runtime (**Fig. 3, Table S4**). The speed of the reference building process is comparable to that of running *de novo* integration since they both start with expression data and require a full pipeline of scaling, PCA, and Harmony integration. However, a reference need only be built and saved once in order to map all subsequent query datasets onto it. For instance, initially building a 500,000-cell reference with Symphony took 5,163 seconds (86.1 min) and mapping a subsequent 10,000-cell query onto it took only 0.99 secs, compared to 4,806 secs (80.1 mins) for *de novo* integration on all cells. Symphony offers a 5000x speedup in this application. These results show that Symphony scales efficiently to map against multimillion-cell references, enabling it to power potential web-based queries within seconds.

**Figure 3.**
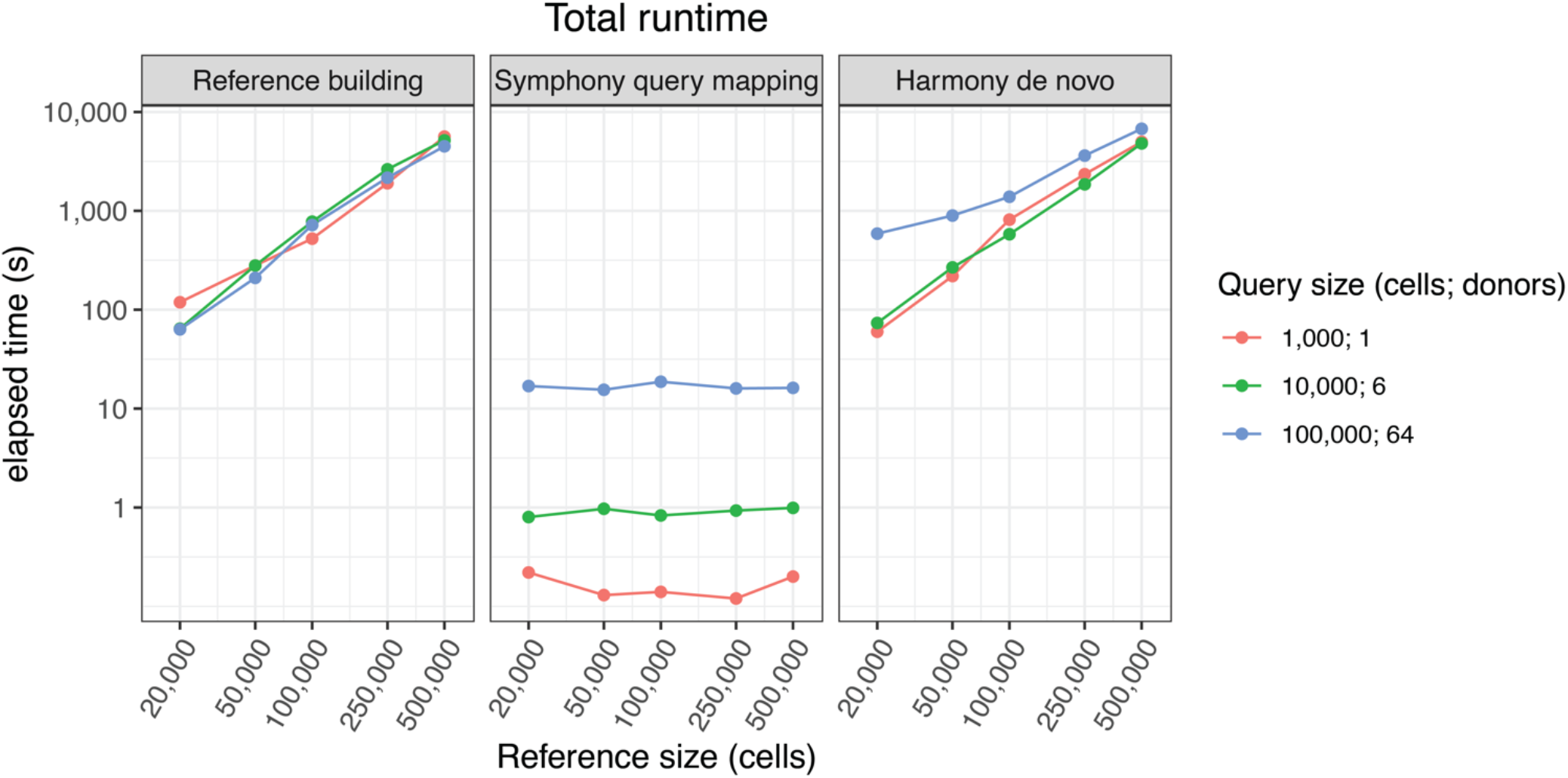
Symphony scales mapping to large references within seconds. Total elapsed time (in secs) required to run Symphony reference building starting from gene expression (left), Symphony query mapping starting from query gene expression (middle), or *de novo* Harmony integration (right) for different-sized reference (x-axis) and query (colors) datasets downsampled from the memory T cell CITE-seq dataset. See Table S4.

Importantly, Symphony mapping time does not depend on the number of cells or batches in the reference since the reference cells are modeled post-batch correction (**Methods**); however, it does depend on the reference complexity (number of centroids *k* and dimensions *d*) and number of query cells and batches (**Table S4**) since the query mapping algorithm solves for the query batch coefficients for each of the reference-defined clusters.

### Symphony maps multi-donor, multi-species study to reference of human pancreatic islet cells

A query dataset might include data from multiple donors, species, and perturbations that create confounding signals obscuring biological signal of interest. Integration algorithms remove these signals in *de novo* analysis, and it is essential that reference mapping removes them too. Therefore, we designed Symphony to simultaneously handle both tasks: mapping query to reference cells and integration within the query. To test the ability of Symphony to integrate query datasets during mapping, we analyzed reference and query datasets of pancreatic islet cells in which both the reference and query have complex experimental structure (**Fig 4a**). The reference contained 5,887 pancreatic islet cells from 32 human donors across four independent studies^41–44^, each profiled with a different plate-based scRNA-seq technology (CEL-seq, CEL-seq2, Smart-seq2, and Fluidigm C1). We manually annotated cell types using cluster-specific marker genes within each reference dataset separately (**Methods**). The query contained 8,569 pancreatic islet cells from 4 human donors and 1,866 cells from 2 mice, all profiled with inDrop, a droplet-based scRNA-seq technology absent in the reference^45^ (**Fig 4b**). PCA of the query dataset alone demonstrated the magnitude of the confounding species and donor signals, emphasizing the need for within-query integration (**Fig. S4a**).

**Figure 4.**
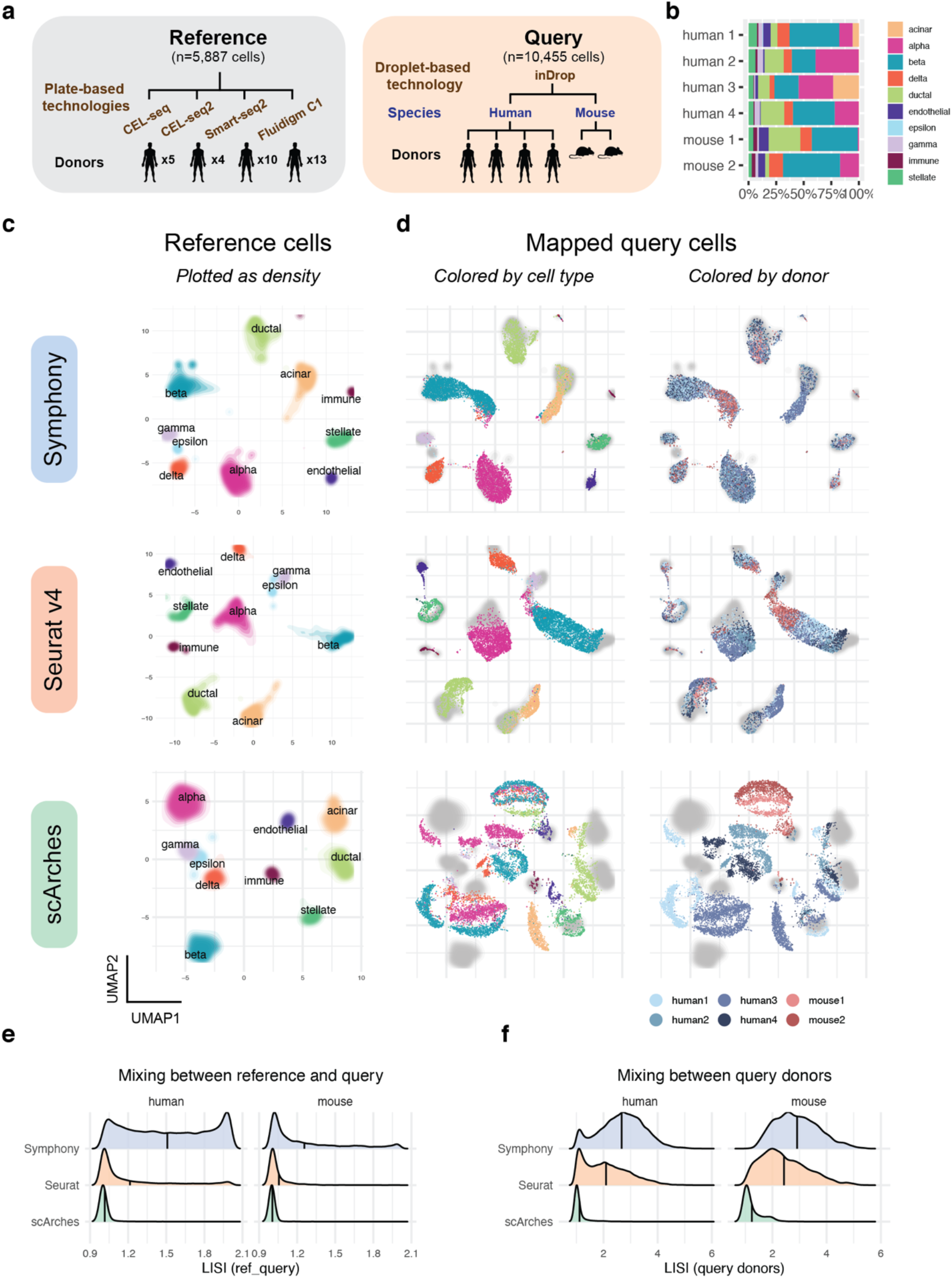
Symphony maps multi-donor, multi-species study to human pancreatic islet cell reference. **(a)** Schematic of mapping experiment with reference (n=5,887 cells, 32 donors) built from four human pancreas datasets and query dataset (n=10,455 cells, from 4 human donors and 2 mouse donors) sequenced on a new technology (inDrop). **(b)** Bar plot shows relative proportions of cell types per query donor. We integrated the reference datasets *de novo* using Harmony, Seurat anchor-based integration, or trVAE, then mapped the query onto the corresponding reference using Symphony, Seurat v4, or scArches, respectively. UMAP plots of the resulting joint embeddings showing **(c)** density of integrated reference cells colored by cell type and **(d)** query cells colored by cell type as defined by Baron et al. (left) or donor identity (right) with reference densities plotted in the back in gray. Degree of integration for each method was measured by LISI metric between reference and query labels **(e)** and LISI between query donors **(f)** for each query cell neighborhood.Distributions of LISI scores for each method faceted by species and normalized to equal height. See Fig.S4 and S5.

Symphony mapped the multi-species, multi-donor, droplet-based query into the reference by effectively and simultaneously removing the effects of species, donor, and technology (**Fig. 4c-d**); reference mapping obtained superior integration compared to PCA (mean donor LISI=2.72 compared to 1.45). We predicted that integrating over three nested sources of variation would make it possible to accurately predict query cell types. Using a simple 5-NN classifier in the harmonized embedding, we observed accurate cell-type prediction. Using ground truth labels defined by the original publication^45^, we obtained a median cell type F1-score of 0.96 (overall accuracy 96%) for human and median cell type F1 of 0.95 (overall accuracy 91%) for mouse cells (**Fig. S4c-d, Table S5**), By mapping against a reference, Symphony is able to overcome strong species effects and simultaneously map analogous cell types between mouse and human.

Next, we evaluated the ability of the other reference mapping algorithms, scArches and Seurat v4, to integrate the same query dataset. For each mapping method, we built a reference using its compatible *de novo* integration method (**Methods, Fig. 4c, S4b**). Symphony obtained higher levels of integration than did Seurat and scArches, both between reference and query as well as donors within the query (**Fig. 4e-f**). Symphony mapping achieves comparable donor mixing to that of Harmony *de novo* integration of all five datasets (mean mapping LISI=2.67 vs *de novo* LISI=2.55 in human, 2.91 vs 2.7 in mouse). In contrast, the other mapping methods return less integrated embeddings, when compared to their corresponding *de novo* methods (mean mapping LISI=2.09 vs *de novo* LISI=2.83 for Seurat in human, 2.43 vs 2.67 in mouse; 1.12 vs 2.52 for scArches/trVAE in human, and 1.24 vs 3.05 in mouse; **Table S6**). We then evaluated the accuracy of each mapping with 5-NN cell type classification (**Methods**). We observed that Symphony and Seurat performed comparably well, and both outperformed scArches on both human and mouse cell type prediction (**Fig. S4c-d, Table S5**). Symphony was 1-2 orders of magnitude faster (1.4 s) than either Seurat (31.7 s) or scArches (381.5 s) mapping on this example (**Table S6**).

### Localizing query cells along a reference-defined trajectory of human fetal liver hematopoiesis

A successful mapping method should position cells not only within cell type clusters but also along smooth transcriptional gradients, commonly used to model differentiation and activation processes over time (**Fig. 5a**). To test Symphony in a gradient mapping context, we built and mapped to a reference atlas profiling human fetal liver hematopoiesis, containing 113,063 liver cells from 14 donors spanning 7-17 post-conceptional weeks of age and 27 author-defined cell types, sequenced with 10x 3’ chemistry (**Fig. 5b, Fig. S6a**)^46^. Trajectory analysis of immune populations with the force directed graph (FDG) algorithm^46^ highlights relationships among progenitor and differentiated cell types (**Fig. 5c**).Notably, the hematopoietic stem cell and multipotent progenitor population branches into three major trajectories, representing the lymphoid, myeloid, and megakaryocyte-erythroid-mast (MEM) lineages. This reference contains two forms of annotation for downstream query inference: discrete cell types and positions along differentiation gradients.

**Figure 5.**
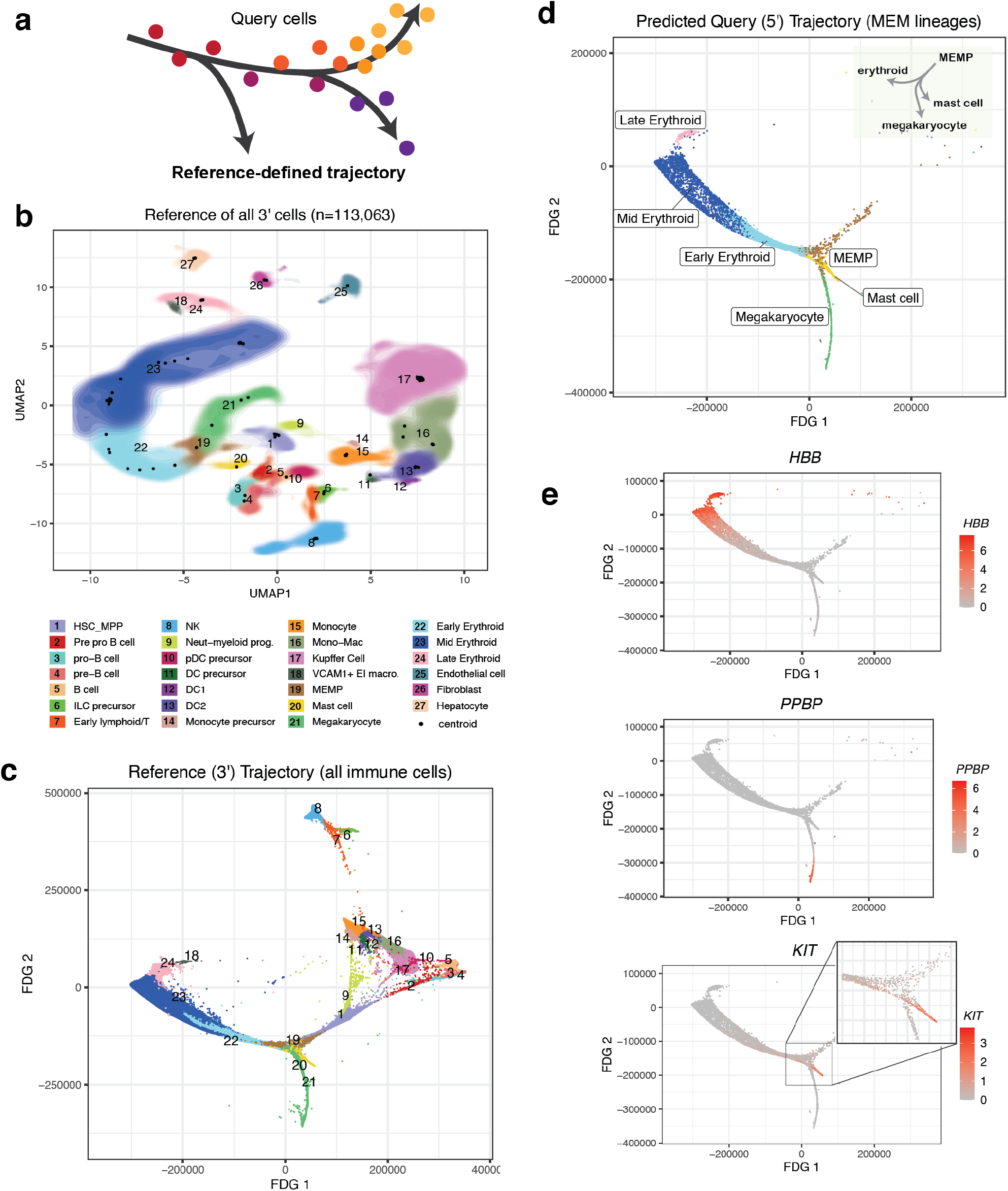
Localizing query cells along a trajectory of fetal liver hematopoiesis. **(a)** Symphony can precisely place query cells along a reference-defined trajectory. The reference (n=113,063 cells, 14 donors) was sequenced using 10x 3’ chemistry, and the query (n=25,367 cells, 5 donors) was sequenced with 10x 5’ chemistry. **(b)** Symphony reference colored by cell types as defined by Popescu et al. (2019). Contour fill represents density of cells. Black points represent soft-cluster centroids in the Symphony mixture model. **(c)** Reference developmental trajectory of 3’-sequenced immune cells (FDG coordinates obtained from original authors). Query cells in the MEM lineages (n=5,141 cells) were mapped against the reference and query coordinates along the trajectory were predicted with 10-NN **(d)**. The inferred query trajectory preserves branching within the MEM lineages, placing terminally differentiated states on the ends. **(e)** Expression of lineage marker genes (*PPBP* for megakaryocytes, *HBB* for erythroid cells, and *KIT* for mast cells). Cells colored by log-normalized expression of gene. See Fig. S6 and S7.

We mapped a query consisting of 21,414 new cells from 5 of the original 14 donors, sequenced with 10x 5’ chemistry. We first inferred query cell types with k-NN classification (**Methods**) and confirmed accurate cell type assignment based on the authors’ independent query annotations^46^ (median cell type F1=0.92 across 14 held-out donor experiments within 3’ dataset only, median cell type F1=0.83 for the 5’-to-3’ experiment; **Fig. S7, Table S7**). To evaluate query trajectory inference, we used the Symphony joint embedding to position query cells from the MEM lineage (n=5,141) in the reference-defined trajectory by averaging the 10 nearest reference cell FDG coordinates. The inferred query trajectory (**Fig. 5d**) recapitulated known branching from MEM progenitors (MEMPs, brown) into distinct megakaryocyte (green), erythroid (blue, pink), and mast cell (yellow) lineages. Moreover, transitions from MEMPs to differentiated types were marked by gradual changes in canonical marker genes (**Fig. 5e**): *PPBP* for megakaryocytes, *HBB* for erythrocytes, and *KIT* for mast cells. These gradual expression patterns are consistent with correct placement of query cells along differentiation gradients.

### Inferring query surface protein marker expression by mapping to a reference assayed with CITE-seq

Recent technological advances in multimodal single-cell technologies (e.g., CITE-seq) make it possible to simultaneously measure mRNA and surface protein expression from the same cells using oligonucleotide-tagged antibodies^47,48^. With Symphony, we can construct a reference from these data, map query cells from experiments that measure only mRNA expression, and infer surface protein expression for the query cells to expand possible analyses and interpretations (**Fig. 6a**).

**Figure 6.**
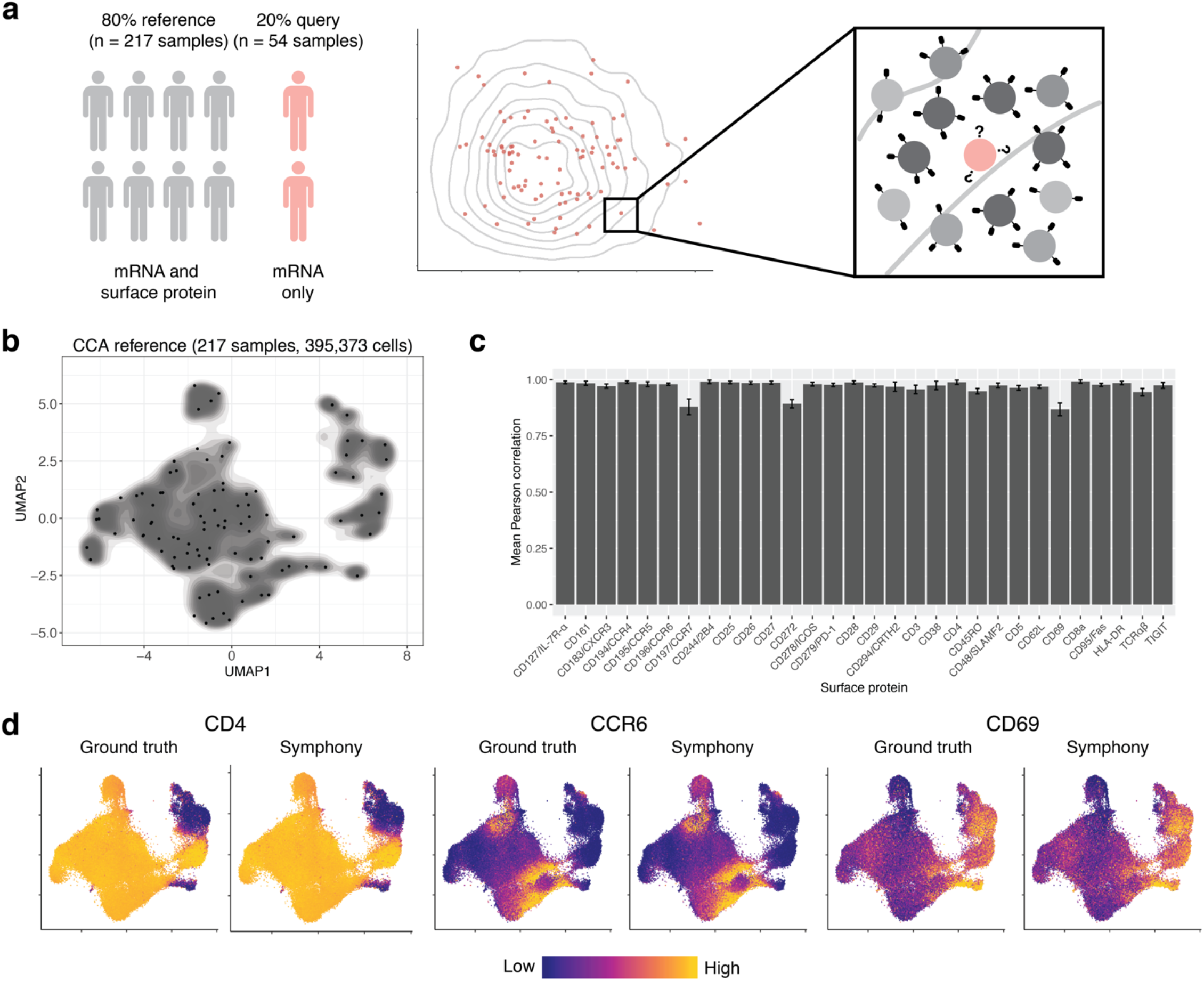
Mapping onto a multimodal reference to infer query surface protein expression in memory T cells. **(a)** Schematic of multimodal mapping experiment. The dataset was divided into training and test sets (80% and 20% of samples, respectively). The training set was used to build a Symphony reference, and the test set was mapped onto the reference to predict surface protein expression in query cells (pink) based on 50-NN reference cells (gray). **(b)** Symphony reference built from mRNA/protein CCA embedding. Contour fill represents density of reference cells. Black points represent soft-cluster centroids in the Symphony mixture model. **(c)** We measured the accuracy of protein expression prediction with the Pearson correlation between predicted and ground truth expression for each surface protein across query cells in each donor. Bar height represents the average per-donor correlation for each protein, and error bars represent standard deviation. **(d)** Ground truth and predicted expression of CD4, CCR6, and CD69 based on CCA reference. Ground truth is the 50-NN-smoothed expression measured in the CITE-seq experiment. Colors are scaled independently for each marker from minimum (blue) to maximum (yellow) expression. See Fig. S8.

To demonstrate this, we used a CITE-seq dataset that measures the expression of whole-transcriptome mRNA and 30 surface proteins on 500,089 peripheral blood memory T cells from 271 samples^40^. We leveraged both mRNA and protein features to build a multimodal reference from 80% of samples (n=217) and map the remaining 20% of samples (n=54). Instead of using PCA, which is best for one modality^49^, we used canonical correlation analysis (CCA) to embed reference cells into a space that leverages both. Specifically, CCA constructs a pair of correlated low-dimensional embeddings, one for mRNA and one for protein features, each with a linear projection function akin to gene loadings in PCA. We corrected reference batch effects in CCA space with Harmony and built a Symphony reference (**Fig. 6b**), saving the gene loadings for the CCA embedding from mRNA features. Then, we mapped the held-out query using only mRNA expression to mimic a unimodal scRNA-seq experiment, reserving the measured query protein expression as a ground truth for validation. We accurately predicted the surface protein expression of each query cell using the 50-NN average from the reference cells in the harmonized embedding. For all proteins, we found strong concordance between predicted and (50-NN smoothed) measured expression (Pearson r: 0.88-0.99, **Fig. 6c-d**). For all but three proteins, we achieved comparable results with as few as 5 or 10 nearest neighbors (**Fig. S8a**).

We note that it is also possible to conduct the same analysis with a unimodal PCA-based reference built from the cells’ mRNA expression only. This approach has slightly worse performance for some proteins (Pearson r: 0.65-0.97, **Fig. S8b-d**), demonstrating that a reference built jointly on both mRNA and protein permits better inference of protein expression than an mRNA-only reference, which is consistent with previous observations that mRNA expression is not fully representative of protein expression^47,48^. This analysis highlights how users can start with a low-dimensional embedding other than PCA, such as CCA, to better capture rich multimodal information in the reference.

## Discussion

Mapping query cells into large, annotated references in real time and without the need to share sensitive information from the reference datasets is becoming increasingly important for reproducible single-cell analysis. We approached this inherently complex, big-data problem using well-established mathematical methods from integration analysis. We framed reference mapping as a specialized case of integration between one relatively small dataset and a second larger, more comprehensive, and previously integrated dataset. Because the reference is already integrated, it is natural to use the same mathematical framework from the integration to perform mapping. For instance, the scArches^28^ algorithm uses an autoencoder-based framework to map to references built with autoencoder-based integration algorithms^32,33^. Similarly, Symphony uses the mixture modeling framework to map to references built with Harmony mixture modeling integration. Symphony compresses the reference by extracting relevant reference-derived parameters from the mixture model to map query cells in seconds. With this compression, references can be distributed without the need to share raw expression data or donor-level metadata, which enables data privacy^50^. Symphony compression greatly reduces the size of a reference dataset: for the memory T cell dataset of 500,089 cells, the raw expression matrix is 8.9 GB, whereas the Symphony minimal reference elements are 1.3 MB.

Useful reference atlases contain annotations absent in the query, such as cell type labels (**Fig. 4**), trajectory coordinates (**Fig. 5**), or multimodal measurements (**Fig. 6**). Transfer of these annotations from reference to query is an open area of research that includes algorithms for automated cell type classification^31,35–38^. We approach annotation transfer in two steps. We first learn a predictive model in the reference embedding, and then map query cells and use their reference coordinates to predict query annotations. In this two-step approach, Symphony mapping provides a feature space but is otherwise independent from the choice of downstream inference model. In PBMC type prediction (**Fig. S3**), we used Symphony embeddings to train multiple competitive classifiers: k-NN, SVM, and logistic regression. In our analyses, we were encouraged to find that a simple k-NN classifier can achieve high performance with only 5-10 neighbors. In practice, users can choose more complex inference models if it is warranted for certain annotation types. Moreover, we expect prediction results to improve with more accurate and reproducible annotation methods, such as consistent cell type taxonomies provided by the Cell Ontology^51^ project and better modeling of multimodal expression data^52^.

Because mapping is a special case of integration, we expected Symphony mapping to recapitulate the results of *de novo* Harmony integration. To this end, we defined three conditions under which Symphony and *de novo* integration with Harmony yield equivalent results. In subsequent examples, we showed that Symphony still performs well when the last two conditions are relaxed. The pancreas query contains more cells than its reference (**condition II**), while the liver hematopoiesis reference and query overlap in donors (**condition III**). Condition I, which requires comprehensive cell type coverage in the reference, is less flexible. When the query contains a brand new cell type, it will be aligned to its most transcriptionally similar reference cluster. Note that condition I only pertains to cell types and not clinical and biological contexts. For instance, we successfully mapped mouse pancreas query to an entirely human pancreas reference (**Fig. 4**), because the same pancreatic cell types are shared in both species. Mapping novel cell types is a current limitation and important direction for future work. For now, we advise users interested in novel cell type discovery to supplement a Symphony analysis with *de novo* analyses of the query alone.

Instead of one monolithic reference for all cell types across all tissues and disease, we expect the proliferation of multiple, well-annotated specialized references that focus on fine-grained modeling of diverse biological systems. For instance, the memory T cell reference (**Fig. 6**) will be useful to annotate fine-grained T cell states, while an unsorted PBMC reference (**Fig. 2**) would better suit coarse-grained annotation of multiple immune populations. Similarly, a reference with only healthy individuals is useful for annotation of cell types, while a reference with both healthy and diseased individuals is useful for annotation of cell types and pathological cell states. We advise Symphony users to carefully select the appropriate reference atlas for their study and potentially map to multiple references, as needed. For instance, one may use a PBMC reference to identify and isolate T cells and a memory T cell reference to assign fine-grained labels to query T cells.

As large-scale tissue and whole-organism single-cell reference atlases become available in the near future, Symphony will enable investigators to leverage the rich information in these references to perform integrative analyses and transfer reference coordinates and diverse annotations to new datasets in a rapid and reproducible manner.

## Supporting information

Supplementary Equations

## Acknowledgements

We thank members of the Raychaudhuri Lab for helpful feedback and comments. We thank members of the Tuberculosis Research Unit (TBRU) LIMAA and Socios En Salud, in particular Megan Murray, Jessica Beynor, Yuriy Baglaenko, Sara Suliman, Ildiko van Rhijn, and Leonid Lecca, for their contributions to generating the memory T cell dataset. We would also like to thank Issac Goh, Muzlifah Haniffa, and other members of the Haniffa Lab for graciously providing preprocessed datasets from their fetal liver hematopoiesis study. This work is supported in part by funding from the National Institutes of Health (1UH2AR067677, U19 AI111224, U01 HG009379, 1R01AR073833, and R01AR063759). The project described was supported by award Number T32GM007753 from the National Institute of General Medical Sciences (JBK). The content is solely the responsibility of the authors and does not necessarily represent the official views of the National Institute of General Medical Sciences or the National Institutes of Health.

## Author contributions

I.K., J.B.K., and S.R. conceived the project. J.B.K. and I.K. developed the method and performed the analyses under the guidance of S.R. F.Z. assisted with benchmarking. S.R., A.N., and D.B.M. contributed to generating the memory T cell dataset. A.N. performed analysis of the memory T cell dataset.All authors participated in interpretation and writing the manuscript.

## Figure Legends

**Supplementary Figure 1. Overview of reference mapping pipeline and Symphony data structures. (a)** The overall analysis pipeline comprises various functions (orange boxes) that each perform a transformation on the data. Symphony mapping takes in a query gene expression matrix and a Symphony reference built from integrated reference datasets, and outputs the query cell locations in the harmonized feature embedding. Models trained on the reference feature embedding (e.g. cell type classifier) can transfer annotations to the query for various downstream tasks. **(b)** Steps of reference building algorithm. Reference datasets spanning multiple batches are aggregated into a single expression matrix on which PCA and Harmony integration is performed. The output of reference compression is the Symphony minimal reference elements, consisting of data structures *μ,σ, U, Y*_*cos*_, *N*_*r*_, and *C* (red symbols). Z_r_corr_ (the harmonized reference embedding) is not used for the mapping calculation but is saved for downstream annotation transfer. **(c)** Steps of query mapping algorithm, indicating where each reference element is used. Query cells are projected into reference PCA space, clustered to reference centroids, and corrected to harmonized space by removing query batch effects.

**Supplementary Figure 2. Nearest neighbor correlation (k-NN-corr) metric**. The k-NN-correlation metric assesses how well an alternative embedding recapitulates the structure of a gold standard embedding. k-NN-corr is asymmetric in that it matters which of the two embeddings is selected as the gold standard. Consider a gold standard embedding **(a)** and two alternative embeddings **(b)** and **(c)**, representing a good mapping and a bad mapping, respectively. For a given query cell *q* (red), we identify its top *k* nearest reference (gray) neighbors in the gold standard embedding (*k* = 3 depicted) and calculate the similarity between the query cell and each neighbor. The similarities between the same query-reference neighbor pairs are then calculated in the alternate embedding. k-NN-corr is the Spearman correlation between the similarities in the gold standard vs. alternative embedding, ranging from -1 to +1. Example k-NN-corr for one query cell and *k* = 500 for the **(d)** Symphony embedding and **(e)** PCA projection embedding. **(f)** k-NN-corr distribution across query cells for k=500 and a gold standard Harmony embedding, for either the Symphony embeddings (blue) or a simple PCA projection with no correction step (red), faceted by query dataset.

**Supplementary Figure 3. Symphony performance against automatic cell type classifiers**. Following the cross-technology PBMC benchmarking experiment from Abdelaal et al. (2019), we ran a total of 48 train-test experiments per Symphony-based classifier. Two different versions of the Symphony feature embeddings were generated depending on variable gene selection method: top 2000 variable genes (vargenes) or top 20 differentially genes (DEGs) expressed per cell type. Symphony embeddings were used to train 3 downstream classifiers: k-NN (k=5), SVM with radial kernel, and multinomial logistic regression (glmnet) with ridge. **(a)** Symphony (orange) median cell-type F1 score across 48 train-test experiments compared to supervised methods (green), demonstrating noninferiority to the top supervised methods and stable performance regardless of downstream classification method. Red dot indicates mean of median F1 scores across 48 experiments (used for ordering the methods along the x-axis). **(b, c)** Median cell type F1 score across 48 experiments for the 5-NN classifier with variable gene selection **(b)** and DEG selection **(c)**. Non-diagonal values represent train on one technology, test on another (42 experiments, all with donor 1). Values along the diagonal indicate train on donor 1, test on donor 2 of the same technology (6 experiments; missing square because donor 2 not sequenced with 10x v3).

**Supplementary Figure 4. Comparison of Symphony to alternative reference mapping methods on a cross-species pancreatic islet cell benchmark. (a)** Standard PCA pipeline applied to the Baron et al. query dataset exhibits strong species and donor effects, demonstrating the need for within-query integration.We benchmarked Symphony mapping (on a Harmony-integrated reference), Seurat v4 mapping (on a Seurat anchor-based-integrated reference), and scArches mapping (on a trVAE-integrated reference). For each approach, we built an integrated reference **(b)**, mapped the query, then predicted query cell types using a 5-NN classifier to transfer annotations using the respective reference embedding. **(c)** Query cell prediction accuracy by species for each method as measured by cell type F1 score, with author-defined ground truth labels. Mouse samples did not have acinar or epsilon cells. The resulting joint cell embedding for each tool was visualized by UMAP **(b, d)**: **(b)** Reference cells colored by dataset/technology. **(d)** Query cells colored by correct (green) or incorrect (red) cell type prediction.

**Supplementary Figure 5. Comparison of *de novo* integration methods for harmonizing all five pancreatic islet cell datasets**.As a comparison to reference mapping (Fig 3), we integrated all five pancreatic islet cell technologies (n=16,342 cells) using 3 *de novo* integration methods: Harmony,Seurat anchor-based integration, and trVAE.UMAP visualizations for the integrated embedding colored by batch **(a)** and cell types **(b)** for each method. Cell types for reference datasets (c1, celseq, celseq2, smartseq) were defined within each dataset separately based on marker genes. Query cell types were defined by Baron et al. Degree of mixing between reference and query datasets **(c)** and mixing between query donors **(d)** was measured with LISI metric on query cell neighborhoods for each method, demonstrating equivalent mixing among *de novo* integration methods (compare to Fig 3d-e).

**Supplementary Figure 6. Mapping to a fetal liver hematopoiesis trajectory. (a)** Size and cell type composition of each donor sample in the 10x 3’ dataset across 27 author-defined cell types from Popescu et al. (2019). pcw = post-conception weeks. **(b)** Library complexity for each sample in 10x 3’ and 10x 5’ datasets, showing low complexity for donor F2 and F5 5’-sequenced samples (removed from further analysis). **(c)** UMAP projections of query cells into reference UMAP space after Symphony mapping, faceted by query donor, colored by cell type. Reference UMAP embedding in bottom-right.

**Supplementary Figure 7. Fetal liver hematopoiesis cell type classification confusion matrices**. We performed two versions of the reference mapping experiments to assess cell type classification accuracy across 27 fine-grained cell types: (1) using exclusively 10x 3’ data, we mapped one held-out donor against a reference constructed from the remaining 13 donors (total 14 mapping experiments), (2) mapping all 10x 5’ cells against all 10x 3’ cells. Cell type confusion matrices are shown for a 30-NN cell type classifier **(a)** aggregated across the 14 held-out donor experiments using exclusively 3’ data and **(b)** the 5’-to-3’ experiment mapping the full 5’ query (n=21,414, n=5 donors) against the full 3’ reference (n=113,063 cells, 14 donors), colored by the proportion of the true cell type that was classified correctly. True cell type is defined by the original authors (Popescu et al., 2019). Rows (true query cell types) are sorted by hierarchical clustering on the average gene expression (all genes) for the cell types to order similar types together. Bar graph (right) shows population size for each cell type.

**Supplementary Figure 8. Inferring query surface protein expression in memory T cells. (a)** Mean Pearson correlation for CCA reference between k-NN predicted protein expression and ground truth for different values of *k*. **(b)** Symphony reference built from a standard mRNA PCA embedding (reference protein values were not used to build embedding but treated as annotations only). Contour fill represents density of reference cells. Black points represent soft-cluster centroids in the Symphony mixture model. **(c)** We measured the accuracy of protein expression prediction based on the PCA reference with the Pearson correlation between predicted and ground truth expression for each surface protein across query cells in each donor. Bar height represents the average per-donor correlation for each protein, and error bars represent standard deviation. **(d)** Ground truth and predicted expression of CD4, CCR6, and CD69 based on PCA reference. Ground truth is the 50-NN-smoothed expression measured in the CITE-seq experiment. Colors are scaled independently for each marker from minimum (blue) to maximum (yellow) expression.

**Supplementary Table 1**. Links to datasets used in the study.

**Supplementary Table 2**. Canonical lineage markers (Wilcoxon rank sum test and auROC statistic) and top 10 differentially expressed genes per cluster used to assign cell types in 10x PBMCs.

**Supplementary Table 3**. Cell type classification confusion matrices for the three 10x PBMCs mapping experiments.

**Supplementary Table 4**. Runtime scalability analysis results (downsampling memory T cell dataset), showing effect of reference and query size, number of query cells or donors, and number of reference centroids or embedding dimensions on elapsed time (in secs).

**Supplementary Table 5**. Cell type classification confusion matrix for multi-donor, multi-species pancreatic islet cell benchmarking example (mapping Baron et al. 2016 as query) among the reference mapping methods evaluated.

**Supplementary Table 6**. Degree of mixing between reference and query cells (ref_query LISI) and between donors within the query (query donor LISI) as well as runtime comparison across different reference mapping methods and corresponding *de novo* integration methods (Symphony/Harmony, Seurat v4/Seurat, and trVAE/scArches) for multi-donor, multi-species pancreas benchmarking example.

**Supplementary Table 7**. Cell type classification confusion matrix for mapping 10x 5’-sequenced fetal liver cells onto an atlas of 3’-sequenced fetal liver cells (Popescu et al. 2019). True labels provided by the original authors, and predictions were made using a 30-NN classifier.

## Methods

### 1. Symphony

#### 1.1 Symphony overview

The goal of single-cell reference mapping is to embed newly assayed query cells into an existing comprehensive reference atlas, facilitating the automated transfer of annotations from the reference to the query. The optimal mapping method needs to be able to operate at various levels of resolution, capture continuous intermediate cell states, and scale to multimillion cells^27^. Consider a scenario in which we wish to map a query of *m* cells against reference datasets with *n* cells, where *m<<n*. Unsupervised integration of measurements across donors, studies, and technological platforms is the standard way to compare single cell datasets and identify cell types. Hence, a “gold standard” reference mapping strategy might be to run Harmony integration on all *m+n* cells *de novo*. However, this approach is impractical because it is cumbersome and time-intensive to process all the cell-level data for the reference datasets every time a user wishes to reharmonize it with a query. Instead, we envision a pipeline where a reference atlas need only be carefully constructed and integrated once, and all subsequent queries can be rapidly mapped into the same stable reference embedding. Symphony is a reference mapping method that efficiently places query cells in their precise location within an integrated low-dimensional embedding of reference cells, approximating *de novo* harmonization without the need to reintegrate the reference cells. Symphony is comprised of two algorithms: reference compression and mapping. Expanding upon the linear mixture model framework introduced in Harmony^17^, Symphony compression takes in an integrated reference and faithfully compresses it by capturing the components of the model into efficient data structures. The output of reference compression is the minimal set of elements needed for mapping **(Fig. S1b)**. The Symphony mapping algorithm takes as input a new query dataset as well as minimal reference elements and returns the appropriate locations of the query cells within the integrated embedding **(Fig. S1c)**.

Once a harmonized reference is constructed and compressed using Symphony, subsequent mapping of query cells executes within seconds (**Fig. 3**). Efficient implementations of Symphony are available as part of an R package at https://github.com/immunogenomics/symphony, along with several precomputed references constructed from public scRNA-seq datasets. The following sections introduce the Symphony model, then describes Symphony compression and mapping in terms of the underlying data structures and algorithms. We also provide **Supplementary Equations** containing more detailed derivations for reference compression terms.

##### Glossary

We define all symbols for data structures used in the discussion of Symphony below, including their dimensions and possible values. Dimensions are in terms of the following parameters:

- *n:* the number of reference cells
- *m:* the number of query cells
- *N:* the total number of cells (*n + m*)
- *g:* the number of genes in the reference after any gene selection
- *d:* the dimensionality of the embedding (e.g. PCs). *d* applies to both reference and query.
- *b:* the number of batches in the reference
- *c:* the number of batches in the query
- *k:* the number of clusters in the mixture model for reference integration (representing latent cell states)

### Reference-related symbols

*G*_*r*_ ∈ *ℝ*^*g* × *n*^ Input reference gene expression matrix, prior to scaling.

*G*_*rs*_ ∈ *ℝ*^*g* × *n*^ Scaled reference gene expression matrix.

*X*_*r*_ ∈ {0,1}^*b* × *n*^ One-hot design matrix assigning reference cells (columns) to batches (rows).

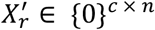 Zero matrix assigning reference cells (columns) to *query* batches (rows). All values are 0 because reference cells do not belong to query batches. This term is used in the derivation for the reference compression terms.

*μ* ∈ *ℝ*^*g* × 1^ Reference gene means used to center each gene for PCA.

*σ* ∈ *ℝ*^*g* × 1^ Reference gene standard deviations used to scale each gene for PCA.

*U* ∈ *ℝ*^*g* × *d*^ Gene loadings from the original PCA (before Harmony integration).

*Z*_*r*_ ∈ *ℝ*^*d* × *n*^ Original (non-harmonized) PC embedding for reference cells.

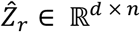 Integrated embedding for reference cells in harmonized PC (hPC) space, as output by Harmony.

*R*_*r*_ ∈ [0,1]^*k* × *n*^ Soft cluster assignment of reference cells (columns) to clusters (rows), as output by Harmony. Each column is a probability distribution that sums to 1.

*Y*_*cos*_ *∈ ℝ*^*d* × *k*^ Cluster centroid locations in the harmonized embedding, L2 normalized.

*B*_*r*_ ∈ *ℝ*^*g* × (1+*b*) × *d*^ 3D tensor of the estimated parameters (betas and intercepts) of the linear mixture model for each of *k* clusters for the reference cells.

*N*_*r*_ ∈ *ℝ*^*k* × *1*^ First reference compression term. Vector containing the size of each of the *k* clusters, effectively the number of reference cells contained within them.

*C* ∈ *ℝ*^*k* × *d*^ Second reference compression term.

*Ref* = {*μ, σ, U, Y*_*cos*,_, *N*_*r*_, *C*} Symphony minimal reference elements comprising *μ, σ, U, Y*_*cos*,_, *N*_*r*_, *C*.

### Query-related symbols

*G*_*q*_ ∈ *ℝ*^*g* × *m*^ Input query gene expression matrix, prior to scaling.

*G*_*qs*_ ∈ *ℝ*^*g* × *m*^ Query gene expression matrix, scaled by *reference* gene means *μ* and standard deviations *σ*

*X*_*r*_ ∈{0,1}^*c* × *m*^ Design matrix assigning query cells (columns) to query batches (rows).

*Z*_*q*_ ∈ *ℝ*^*d* × *m*^ Query cell locations in original (non-harmonized) PC embedding.

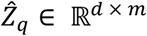 Approximate query cell locations in integrated embedding (hPC space). Output of Symphony reference mapping.

*R*_*q*_ ∈ [0, 1]^= *k* × *m*^ Soft cluster assignment of query cells (columns) to clusters (rows). Each column is a probability distribution that sums to 1.

*B*_*q*_ ∈ *ℝ*^*k* × (1+c) × *d*^ 3D tensor of the estimated parameters (betas and intercepts) of the linear mixture model for each of *k* clusters.

#### 1.2 Symphony model and conditions for equivalence to Harmony integration

Symphony and Harmony both use a linear mixture model framework, but the two methods perform different tasks: Harmony integrates a reference, whereas Symphony compresses the reference and enables efficient query mapping. To motivate the Symphony model, it is helpful to first briefly review the mixture model, which serves as the basis. Harmony integrates scRNA-seq datasets across batches (e.g. multiple donors, technologies, studies) and projects the cells into a harmonized embedding where cells cluster by cell type rather than batch-specific effects. Harmony takes as input a low-dimensional embedding of cells (*Z*) and design matrix with assignments to batches (*X*) and outputs a harmonized embedding 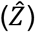 with batch effects removed. Briefly, Harmony works by iterating between two subroutines—maximum diversity clustering and linear mixture model correction—until convergence. In the clustering step, cells are probabilistically assigned to soft clusters with a variant of soft *k-*means with a diversity penalty favoring clusters represented by multiple datasets rather than single datasets. In the correction step, each cluster learns a cluster-specific linear model that explains cell locations in PC space as a function of a cluster-specific intercept and batch membership. Then, cells are corrected by cell-specific linear factors weighted by cluster membership to remove batch-dependent effects. The full algorithm and implementation are detailed in Korsunsky et al. (2019)^17^.

In the scenario of mapping *m* query cells against *n* reference cells, the *de novo* integration strategy would model all cells as in (1), where the *H* subscript denotes the Harmony solution, in contrast to the Symphony model which is presented in (2). Let *X*_*H*_ ∈ {0,1}^(*c*+*b*)×(*m*+*n*)^ represent the one-hot encoded design matrix assigning all cells across batches. 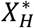 denotes *X*_*H*_ augmented with a row of 1s for the batch-independent intercept term: 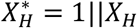 The intercept terms represent cluster centroids (location of “experts” in the mixture of experts model). *Z*_*H*_ represents the low-dimensional PCA embedding of all cells. *R*_*H*_ represents the probabilistic assignment of cells across *k* clusters, and *diag*(*R*_*Hk*=_) ∈ *ℝ*^*N*× *N*^ denotes the diagonalized *k* th row of *R*_*H*_. For each cluster *k*, the parameters of the linear mixture model *B*_*k*=_ ∈ *ℝ*^(1+*c*+*b*) × *d*^ can therefore be solved for as in (1), using ridge regression with ridge penalty hyperparameter *λ*. Note that we do not penalize the batch-independent intercept term: *λ*_*0*_ = 0, ∀ _*a ∈* [1:(*c*+*b*)]_ *λ*_*a*_ = 1.

### *De novo* Harmony model

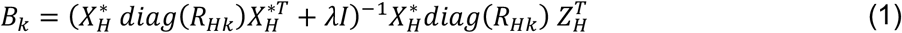

The goal of Symphony mapping is to add new query cells to the model in order to estimate and remove the query batch effects. Symphony mapping approximates *de novo* Harmony integration on all cells, except the reference cell positions in the harmonized embedding do not change. In order for Symphony mapping to be equivalent to *de novo* Harmony, several conditions must be met:

i. All cell states represented in the query dataset are captured by the reference datasets—i.e. there are no completely novel cell types in the query.
ii. The number of reference cells is much larger than the query (*m<<n*).
iii. The query dataset is obtained independent of the reference datasets—i.e. the reference batch design matrix (*X*_*r*_) has no interaction with the query batch design matrix (*X*_*q*_).

We consider these to be fair assumptions for large-scale reference atlases, allowing Symphony to make three key approximations:

1. With a large reference, the reference-only PCs approximate the PCs for the combined reference and query datasets. This allows us to project the query cells into the pre-harmonized reference PCA space using the reference gene loadings (*U*).
2. The cluster centroids (*Y*) for the integrated reference cells approximate the cluster centroids from harmonizing all cells.
3. The reference cell cluster assignments (*R*_*r*_) remains approximately stable with the addition of query cells.

Given these approximations, we can thereby harmonize the reference cells *a priori* and save the reference-dependent portions of the Harmony mixture model (**Supplementary Equations**). In Symphony, we model the reference cells as already harmonized with batch effects removed, so we can thereafter ignore the reference design matrix structure. The Symphony design matrix *X ∈* [0, 1]^C^ ^× *N*^ assigns all cells (reference and query) to *query* batches only. *X*^∗^ denotes *X* augmented with a row of 1s 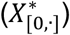 corresponding to the batch-independent intercepts (we model the intercepts for all cells). The remaining *c* rows 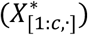 represent the one-hot batch assignment of the cells among the *c* query batches. Note that for the reference cell columns, these values are all 0 since the reference cells do not belong to any *query* batches. The parameters (*B*_*qk*=_ *∈ ℝ* ^(1+c) × *d*^) of the model for each cluster *k* can then be solved for as in (2). Similar to Harmony, we use ridge regression penalizing the non-intercept terms, where *λ*_*0*_ = 0, *∀*_*a∈* [1*:c*]_ *λ*_*a*_ = 1.

### Symphony model

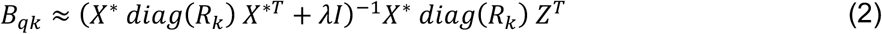

The matrix *R ∈ ℝ* ^*k*×*N*^ denotes the assignment of query and reference cells (columns) across the reference clusters (rows). *Z ∈ ℝ*^*d*×*N*^ denotes the horizontal matrix concatenation of the uncorrected query cells in original PC space (*Z*_*q*_) and corrected reference cells in harmonized space 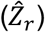 For each cluster *k*, let matrix *B*_*qk*=_ ∈ *ℝ*^(1+*C*) × *d*^ represent the query parameters to be estimated. The first row of *B*_*qk*_ represents the batch-independent intercept terms, and the remaining *c* rows of *B*_*qk*_ represent the query batch-dependent coefficients, which can be regressed out to harmonize the query cells with the reference. Note that the intercept terms from Symphony mapping should equal the cluster centroid locations from the integrated reference since the harmonized reference cells are modeled only by a weighted average of the centroid locations for the clusters over which it belongs (and a cell-specific residual). Hence, the reference cell positions should not change when removing query batch effects.

The matrices *X*^∗^, *R*_*k*_, and *Z* in (2) can be partitioned into query and reference-dependent portions. In the **Supplementary Equations**, we show in detail how the reference-dependent portions can be further simplified into a *k* x 1 vector and *k* x *d* matrix (*N*_*r*_ and C), which we call “reference compression terms.” Intuitively, the vector *N*_*r*_ contains the size (in cells) of each reference cluster. The matrix 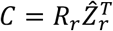 does not have as intuitive an explanation but follows from the derivation (**Supplementary Equations**). These terms can be computed at the time of reference building and saved as part of the minimal reference elements to reduce the necessary computations during mapping.

#### 1.3 Reference building and compression

Reference compression is the key idea that allows for the efficient mapping of new query cells onto the harmonized reference embedding without the need to reintegrate all cells. To construct a Symphony reference with minimal elements needed for mapping, reference cells are first harmonized in a low-dimensional space (e.g. PCs) to remove batch-dependent effects. Symphony then compresses the Harmony mixture model components to be saved for subsequent query mapping.

### Data structures

Symphony takes as input a gene expression matrix for reference cells (*G*_*r*_) and corresponding one-hot-encoded design matrix (*X*_*r*_) containing metadata about assignment of cells to batches. It outputs a set of data structures, referred to as the Symphony minimal reference elements, that captures key information about the reference embedding that can be subsequently used to efficiently map previously unseen query cells (**Algorithm 1**). These components include the gene mean (*μ*) and standard deviation (*σ*) used to scale the genes, the PCA gene loadings (*U*), the final L2-normalized cluster centroid locations (*Y*_*cos*,_), and precomputed values which we call the “reference compression terms” (*N*_*r*_ and *C*) that expedite the correction step of query mapping (**Supplementary Equations**). These elements are a subset of the components available once Harmony integration is applied to the reference cells. Note that other input embeddings, such as canonical correlation analysis (CCA), may be used in place of PCA as long as the gene loadings to perform query projection into those coordinates are saved.

**Table 1** lists the Symphony minimal reference elements required to perform mapping. **Table 2** shows additional components of a “full” Harmony reference that are not included in the Symphony reference elements. Importantly, the dimensions of the Symphony data structures do not require information on the *n* individual reference cells and hence do not scale with the raw number of reference cells. Rather the components scale with the biological complexity captured (i.e. number of clusters *k* and dimensionality of embedding *d*). Conversely, the Harmony data structures store information on a per-cell basis (*n*). Note that in practice the integrated embedding of reference cells 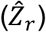 listed in **Table 2** is needed to perform downstream transfer of annotations from reference to query cells (e.g. k-NN), but it is not required during any computations of the mapping step.

**Table 1:** Symphony minimal reference elements.

|  |  |
| --- | --- |
| $\mu \in \mathbb{R}^{g \times 1}$ | Reference gene means used to center each gene for PCA. |
| $\sigma \in \mathbb{R}^{g \times 1}$ | Reference gene standard deviations used to scale each gene for PCA. |
| $U \in \mathbb{R}^{g \times d}$ | Gene loadings to project from expression to PCA (or CCA) space |
| $Y_{cos} \in \mathbb{R}^{d \times k}$ | Cluster centroid locations in harmonized PC space, L2 normalized. |
| $N_r \in \mathbb{R}^{k \times 1}$ | First reference compression term. Vector containing the size of each of the $k$ clusters, effectively the number of reference cells contained within them. |
| $C \in \mathbb{R}^{k \times d}$ | Second reference compression term. |

**Table 2:** Additional components of Harmony reference.

|  |  |
| --- | --- |
| $G_r \in \mathbb{R}^{g \times n}$ | Input reference gene expression matrix, prior to scaling. |
| $X_r \in \{0, 1\}^{b \times n}$ | Design matrix assigning reference cells (columns) to reference batches (rows). |
| $B_r \in \mathbb{R}^{k \times (1+b) \times d}$ | 3D tensor of the estimated parameters (betas and intercepts) of the linear mixture model for each of $k$ clusters for the reference cells. |
| $\hat{Z}_r \in \mathbb{R}^{d \times n}$ | Integrated embedding for reference cells in harmonized PC (“hPC”) space, as output by Harmony. |
| $R_r \in [0, 1]^{k \times n}$ | Soft cluster assignment of reference cells (columns) to clusters (rows), as output by Harmony. Each column is a probability distribution that sums to 1. |

### Algorithm

Starting from reference cell gene expression, we first perform within-cell library size normalization (if not already done) and variable gene selection to obtain *G*_*r*_, scaling of the genes to have mean 0 and variance 1 (saving *μ* and *σ* for each gene), and PCA to embed the reference cells in a low-dimensional space, saving the gene loadings (*U*) (**Implementation Details**). Then, the PCA embedding (*Z*_*r*_) and batch design matrix (*Z*_*r*_) are used as input to Harmony integration to harmonize over batch-dependent sources of variation. Given the resulting harmonized embedding 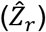 and final soft assignment of reference cells to clusters (*R*_*r*_), the locations of the final reference cluster centroids *Y ∈ ℝ*^*d* × *k*^ can be calculated as in (3) and saved.

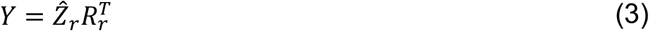

Symphony then computes the reference compression terms *N*_*r*_ (intuitively, the number of cells per cluster) and *C*, which does not have an intuitive explanation but can be directly computed as 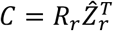 Refer to the **Supplementary Equations** for a complete mathematical derivation of the compression terms. Symphony reference building ultimately returns the minimal reference elements: *μ, σ, U, Y*_*cos*,_, *N*_*r*_, and *C* (**Fig. S1a**).

#### Algorithm 1

Build Symphony reference

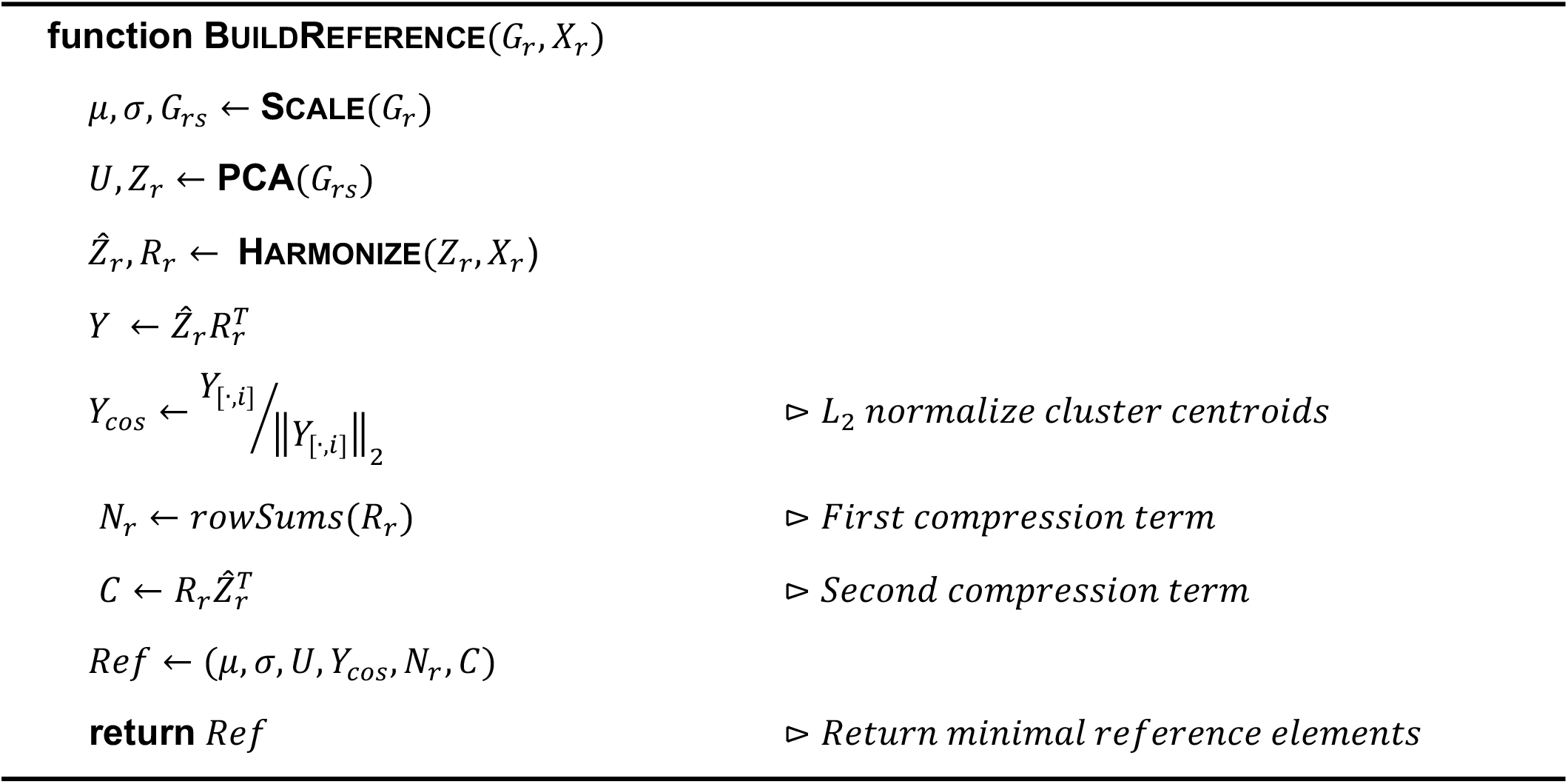

#### 1.4 Symphony mapping

The Symphony mapping algorithm localizes new query cells to their appropriate locations in the harmonized embedding without the need to run integration on the reference and query cells altogether. The joint embedding of reference and query cells can be used for downstream analyses, such as transferring cell type annotations from the reference cells to the query cells.

### Data structures

Symphony mapping takes as input the gene expression matrix for query cells (*G*_*q*_), query design matrix assigning query cells to batches (*X*_*q*_), and the precomputed minimal elements for a reference (*Ref*). It outputs a query object containing the locations of query cells in the integrated reference embedding (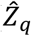 **Algorithm 2**). **Table 3** lists the components of the query object that is returned by Symphony.

**Table 3:** Components of Symphony query.

|  |  |
| --- | --- |
| $G_q \in \mathbb{R}^{g \times m}$ | Input query gene expression matrix, prior to scaling. |
| $X_q \in \{0, 1\}^{c \times m}$ | Design matrix assigning query cells (columns) to query batches (rows). |
| $Z_q \in \mathbb{R}^{d \times m}$ | Query cell locations in original (non-harmonized) PC embedding. |
| $\hat{Z}_q \in \mathbb{R}^{d \times m}$ | Approximate query cell locations in integrated embedding (hPC space). |
| $R_q \in [0, 1]^{k \times m}$ | Soft cluster assignment of query cells (columns) to clusters (rows). Each column is a probability distribution that sums to 1. |
| $B_q \in \mathbb{R}^{k \times (1+c) \times d}$ | 3D tensor of the estimated parameters (betas and intercepts) of the linear mixture model for each of $k$ clusters. |

### Algorithm

The input to the query mapping procedure is a gene expression matrix (*G*_*q*_) and design matrix (*X*_*q*_) for query cells, and the output is the locations of the cells in the harmonized embedding 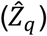. At a high level, the mapping algorithm first projects the query cells into the original, non-harmonized PC space as the reference cells using the reference gene loadings (*U*) and assigns probabilistic cluster membership across the reference cluster centroid locations. Then, the query cells are modeled using the Symphony mixture model and corrected to their approximate locations in the integrated embedding by regressing out the query batch-dependent effects (**Algorithm 2**).

### Projection of query cells into pre-harmonized PC Space

Symphony projects the query cells into the same original PCs (*Z*_*r*_) as the reference. Symphony assumes that, given a much smaller query compared to the reference (*m<<n*), the PCs will remain approximately stable with the addition of query cells. To project the query cells, we first subset the query expression data by the same variable genes used in reference building and scale the normalized expression of each gene by the same mean and standard deviations used to scale the reference cells *(μ, σ*). Let *G*_*qs*,_ denote the query gene expression matrix scaled by the reference gene means and standard deviations. We can then use the reference gene loadings (*U*) to project *G*_*qs*,_ into reference PC space. In (4), *Z*_*q*_ *∈ ℝ*^*d*×*m*^ denotes the PC embedding for the query cells. Note that if an alternate starting embedding (e.g. CCA) is used instead of PCA, the gene loadings must be saved to enable this query projection step.

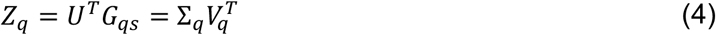

### Soft assignment across reference clusters

Once the query cells are projected into PC space, we soft assign the cells to the reference clusters using the saved reference centroid locations (*Y*_*cos*,_). Symphony assumes that the reference cluster centroid locations remain approximately stable with the addition of a much smaller query dataset since the query contains no novel cell types. Under these conditions, we use a previously published objective function for soft *k*-means clustering (5), which includes a distance term and an entropy regularization term over *R* weighted by hyperparameter *σ*. This is the same objective function as the clustering step of Harmony, except it does not include the diversity penalty term. In Harmony, the purpose of the diversity term is to penalize clusters that are only represented by one or a few datasets (suggesting they do not represent true cell types). In contrast, Symphony does not require the use of a diversity penalty because the reference centroids have already been established. Furthermore, the query cell types can comprise a subset of a larger set of reference cell types, and therefore not all clusters are necessarily expected to be represented in the query. We can solve for *R*_*q*_, the optimal probabilistic assignment for query cells across each of the *k* reference clusters (**Implementation Details**).

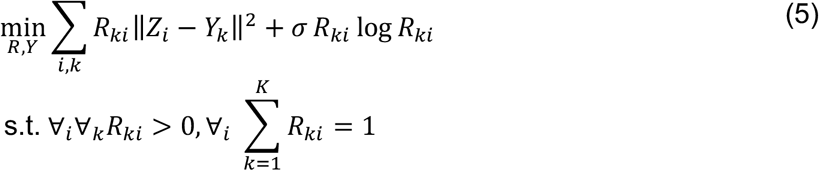

### Mixture of experts correction

The final step in Symphony mapping is to model then remove the query batch effects to obtain 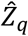, the approximate location of query cells in the harmonized reference embedding. In equation (2), we modeled the reference and query cells together and wish to solve for the query parameters *B*_*qk*_ *∈ ℝ*^(1+*C*) × *d*^ for each cluster *k*. The reference-dependent terms in (2) were previously computed and saved in compressed form (*N*_*r*_ and *C*). With *R*_*q*_ and *Z*_*q*_ calculated from query cell projection and clustering, we can finally solve for *B*_*qk*_. Similar to the correction step of Harmony, we obtain cell-specific correction values for the query cells by removing the batch-dependent terms captured in *B*_*qk*[1*:C*,.]_. Note that the reference batch terms are neither modeled nor corrected during reference mapping, so the harmonized reference cells do not move.

The final locations of the query cells in the harmonized embedding are estimated by iterating over all *k* clusters and subtracting out the non-intercept batch terms for each cell weighted by cluster membership (6). Intuitively, the query centroids are moved so that they overlap perfectly with the reference centroids in the harmonized embedding. 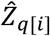 denotes the approximate location in harmonized PC space for query cell *i*.

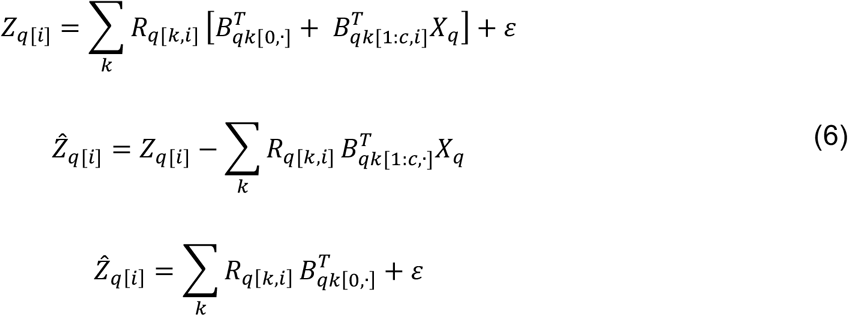

#### Algorithm 2

Map query cells onto reference

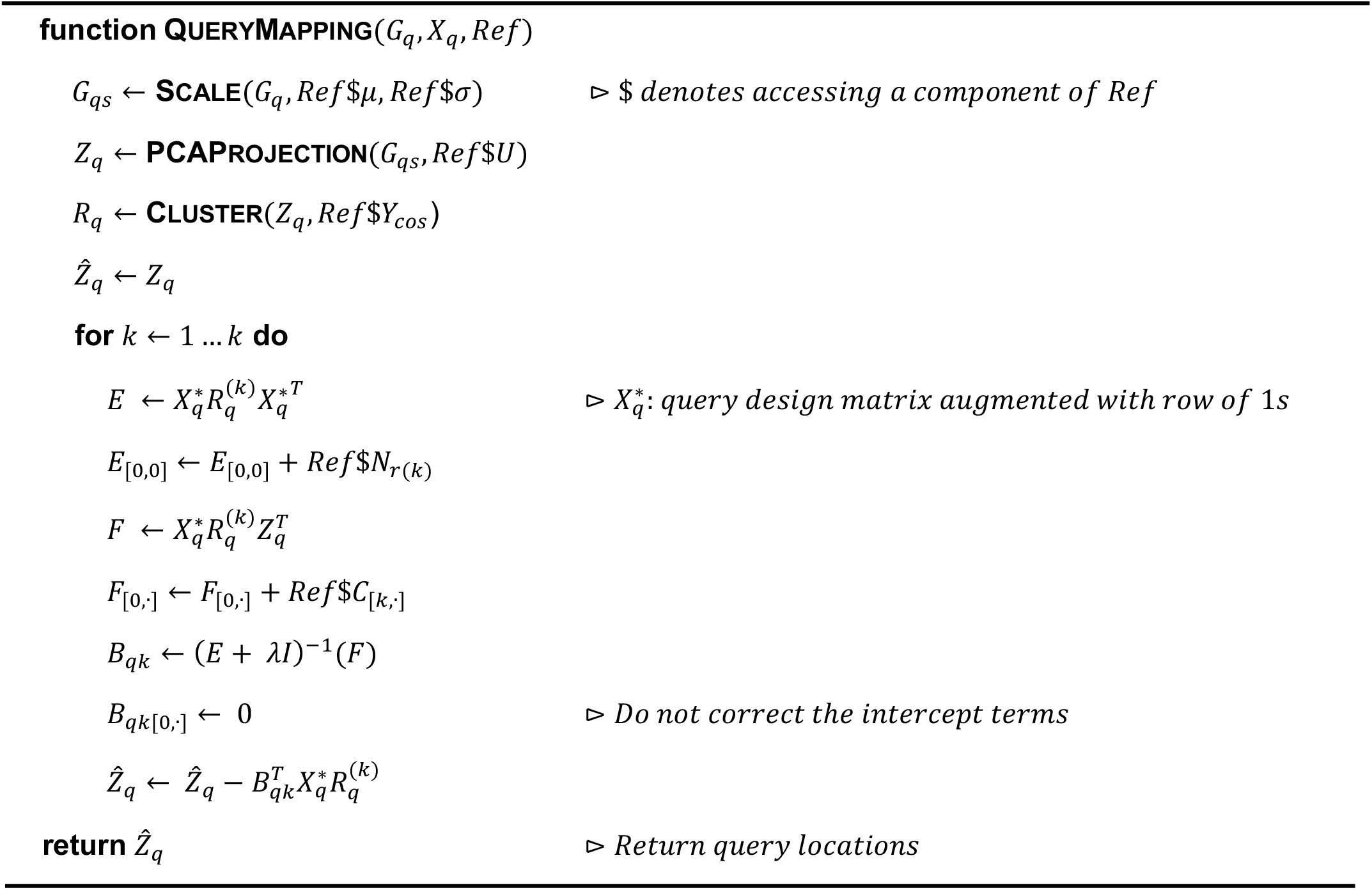

### 1.5 Implementation details

#### Reference building and compression

##### Variable gene selection and scaling

Starting with the gene expression matrix for reference cells, we perform log(CP10K) library size normalization of the cells (if not already done), subset by the top *g* variable genes by the vst method (as provided in Seurat^18^), which fits a line to the log(variance) and log(mean) relationship using local polynomial regression, then standardizes the features by observed mean and expected variance, calculating gene variance on the standardized values, which is re-implemented as a standalone function at https://github.com/immunogenomics/singlecellmethods. The data is scaled such that the expression of each gene has a mean expression of 0 and variance of 1 across all cells.

##### PCA

We perform dimensionality reduction on the scaled gene expression *G*_*rs*,_ using principal component analysis (PCA). PCA projects the data a low-dimensional, orthonormal embedding that retains most of the variation of gene expression in the dataset. Singular value decomposition (SVD) is a matrix factorization method that can calculate the PCs for a dataset. Here, we use SVD (irlba package in R^53^) to perform PCA. SVD states that matrix *G*_*rs*,_ with dimensions *g* × *n* can be factorized as:

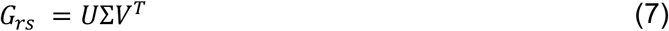

In (7), *∑ v*^*T*^ = *Z*_*r*_ (dimensions *d* × *n*) represents the embedding of reference cells in PC space, after truncating the matrix on the first *d* (by default, *d* = *2*0) PCs. The gene loadings (*U ∈ ℝ*^*g* × *d*^) are saved. Note that an alternative embedding, such as canonical correlation analysis (CCA) may be used in place of PCA, as long as the gene loadings are saved.

##### Harmony integration

The PCA embedding (*Z*_*r*_) is then input to Harmony for dataset integration. By default, Symphony uses the default parameters for the cluster diversity enforcement (*θ* = *2*), the entropy regularization hyperparameter for soft *k*-means (*σ* = 0.1), and the number of clusters 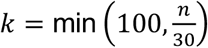. We save the L2-normalized cluster centroid locations *Y*_*cos*,_ to the reference object since query mapping employs a cosine distance metric. If the reference has a single-level batch structure, no integration is performed, and the clusters are defined using soft k-means.

### Query mapping

#### Normalization and scaling

The gene expression for query cells are assumed to be library size normalized in the same manner that was used to normalize the reference cells (e.g. log(CP10K)). During scaling, the query data is subset by the same variable genes from the reference datasets, and query gene expression is scaled by the *reference* gene means and standard deviations. Any genes present in the query but not the reference are ignored, and any genes present in the reference but not the query have scaled expression set to 0.

#### Clustering step uses cosine distance

As in Harmony, in practice we use cosine distance rather than Euclidean distance in the clustering step. For the computation of the distance term, we L2-normalize the columns (cells) of *Z* and columns (centroids) of *Y*_*k*_ such that the squared values sum to 1 across each column. Let the terms *Z*_*q* _ *cos* [·,*i*]_ and *Y*_*cos* [·,*k*]_ represent the L2-normalized locations of query cell *i* and the reference centroid for cluster *k* in PC space, respectively. We compute the cosine distance between the cells and centroids. Since all *Zq*__*cos*_ [.,_*i*_] and *Y* _*cos*_ [.,_*k*_] each have unity norm, the squared Euclidean distance ∥ *Z*_*q*_*cos* [·,*i*]_− *Y*_*cos*_ [·,_*k*_] ∥ ^*2*^ is equivalent to the cosine distance 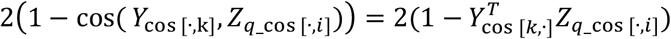 Therefore, the objective function for query assignment to centroids becomes:

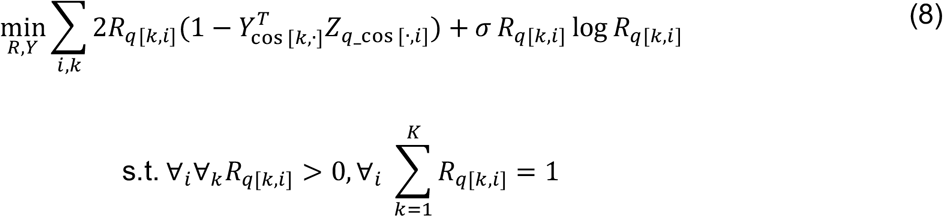

We can solve the optimization problem using an expectation-maximization framework. Following the same strategy as Korsunsky et al. (2019), we calculate *R*_*i*_, the optimal probabilistic assignment for each query cell *i* across each of the *k* reference clusters. In (9), we can interpret *R*_*q*[*k,i*]_ as the probability that query cell *i* belongs to cluster *i*. The denominator term simply ensures that for any given cell *k*, the probabilities across all *i* clusters sum to one. By default, sigma=0.1

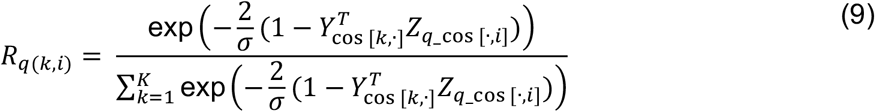

## 2. Analysis details

### 2.1 10x PBMCs analysis

#### Preprocessing scRNA-seq data

The three 10x PBMCs datasets were previously preprocessed by our group as part of the Harmony publication. We used the same log(1+CP10K) normalized expression data, filtered as described in Korsunsky et al. (2019)^17^. The PBMCs consist of cells from three technologies: 3’v1 (n=4,808 cells), 3’v2 (8,372 cells), and 5’ (7,612 cells).

#### Symphony mapping experiments

To construct each of three references for subsequent mapping, we aggregated two reference datasets into a single normalized expression matrix and identified the top 2,000 variable genes across all cells using the variance stabilizing transformation (VST) procedure^18^. We ran Harmony on the top 20 PCs and default 100 clusters, harmonizing over ‘technology’ with default parameters. For Symphony mapping, we specified query ‘technology’ covariate.

#### Constructing gold standard embedding

To construct the gold standard *de novo* Harmony embedding, we concatenated all three datasets together into a single expression matrix, subsetted by the top 2,000 variable genes over all cells, and ran Harmony integration on the top 20 PCs, harmonizing over ‘technology’ with default parameters.

#### Assigning ground truth cell types

We clustered the cells in the gold standard embedding using the Louvain algorithm as implemented in the Seurat functions *BuildSNN* and *RunModularityClustering*^18^. For PBMCs, we used nn_k = 5 (to capture rare HSCs), nn_eps = 0.5, and resolution = 0.8. We labeled clusters with ground truth cell types according to expression of canonical lineage marker genes (**Table S2**). PBMCs were assigned across 7 types: T (*CD3D*), NK (*GNLY*), B (*MS4A1*), Monocytes (*CD14, FCGR3A*), DCs (*FCER1A*), Megakaryocytes (*PPBP*), and HSCs (*CD34*). Clusters were labeled if the AUC (calculated using presto^54^) for the corresponding lineage marker was *Y*0.62. For clusters that did not express a specific lineage marker, we manually assigned a cell type based on the top differentially expressed genes (**Table S2**). PBMCs cluster 20 was identified as low-quality cells (high in mitochondrial genes; **Table S2**). We removed all cells in this cluster (n=94) from further analyses. The final ground truth labels were used in downstream analyses and cell type classification accuracy evaluation.

#### Evaluation of mixing and cell type classification accuracy

To compare dataset mixing between *de novo* integration and mapping, we calculated Local Inverse Simpson Index (LISI) using the *compute_lisi* function from https://github.com/immunogenomics/LISI. For each mapping experiment, we calculated dataset LISI on all cells, then subsetted the results for query cell neighborhoods only to measure the effective number of datasets in the local neighborhood of each query cell.

We predicted query cell types by transferring reference cell type annotations using the *knn* function in the ‘class’ R package (k=5). We calculated overall accuracy across all query cells and cell type F1 scores (the harmonic mean of precision and recall, ranging from 0 to 1). Precision = TP/(TP+FP), recall = TP/(TP+FN), F1 = (2 * precision * recall) / (precision + recall). Cell type F1 was the metric Abdelaal et al. used to benchmark automated cell type classifiers^35^. We used their *evaluate*.*R* script to calculate confusion matrices and F1 scores by cell type.

#### Quantifying local similarity between two embeddings

k-NN-correlation (k-NN-corr) is a new metric that quantifies how well a given alternative embedding preserves the local neighborhood structure with respect to a gold standard embedding. Anchoring on each query cell, we calculate (1) the pairwise similarities to its *k* nearest reference neighbors in the gold standard embedding and (2) the similarities between the same query-reference neighbor pairs in an alternate embedding (**Methods**), then calculate the Spearman (rank-based) correlation between (1) and (2). For similarity, we use the radial basis function kernel: *similarity*(x,y) = exp(-∥x-y∥^2^/(2*σ*^2^)). For each query cell, we obtain a single k-NN-corr value capturing how well the relative similarities to its *k* nearest reference neighbors are preserved. Note that k-NN-corr is asymmetric with respect to which embedding is selected as the gold standard and which is selected as the alternative because the nearest neighbor pairs are fixed based on how they were defined in the gold standard. The distribution of k-NN-corr scores for all query cells can measure the embedding quality, where higher k-NN-corr indicates greater recapitulation of the gold standard. Lower values for *k* assess more local neighborhoods, whereas higher *k* assesses more global structure.

We calculated k-NN-corr between the gold standard Harmony embedding and two alternative embeddings: (1) the full Symphony mapping algorithm (projection, clustering, and correction) and (2) PCA-projection only as a comparison to a batch-naïve mapping. PCA-projection refers to the first step of Symphony mapping, where query cells are projected from gene expression to pre-harmonized PC space: Z_q_ = U^T^G_q_.

### 2.2 Benchmarking against automatic cell type classifiers

We downloaded the PbmcBench benchmarking dataset used by a recent comparison of automatic cell type identification methods^35,39^. For each of 48 train-test experiments previously described^35^, we used the same evaluation metrics (median cell type F1 score) to evaluate Symphony in comparison to the 22 other classifiers. We obtained the numerical F1-score results for the other classifiers for all 48 experiments directly from the authors in order to determine Symphony’s place within the rank ordering of classifier performance.

During reference building, we explored two different gene selection methods: (1) unsupervised (top 2000 variable genes) and (2) supervised based on identifying the top 20 differentially expressed (DE) genes per cell type. Option (2) was included to give Symphony the same information as prior-knowledge classifiers (e.g. SCINA with 20 marker genes per cell type). We used the ‘presto’ package^54^ for DE analysis. No integration was performed because the reference had a single-level batch structure (clusters were simply assigned using soft k*-*means). Onto each of 7 references (each representing 1 protocol for donor pbmc1), we mapped either a second protocol for donor pbmc1 (6 experiments) or the same protocol for donor pbmc2 (1 experiment). Given the resulting Symphony joint feature embeddings, we used three downstream classifiers to predict query cell types: 5-NN, SVM with a radial kernel, and glm_net with ridge^55^. A total of 6 Symphony-based classifiers were tested (2 gene selection methods * 3 downstream classifiers).

### 2.2 Pancreas benchmark

#### Constructing the pancreas query with mouse and human

The pancreas query dataset (Baron et al., 2016; inDrop, n=8,569 human and 1,886 mouse cells) along with author-defined cell type labels were downloaded from https://hemberg-lab.github.io/scRNA.seq.datasets/human/pancreas/. In order to combine the human and mouse matrices into a single aggregated query, we “humanized” the mouse expression matrix by mapping mouse genes to their orthologous human genes. This mapping was computed using the biomaRt R package^56^, `mapping mgi_symbol from the mmusculus_gene_ensembl database to hgnc_symbol from the hsapien_gene_ensembl` database. We added additional ortholog pairs from HomoloGene (https://ftp.ncbi.nih.gov/pub/HomoloGene/build37.2/homologene.data) to obtain a total of 22,578 human to mouse gene ortholog pairs. We represented this map as a matrix, with mouse genes as rows, human genes as columns, and values in {0,1} assigned to denote whether a mouse gene maps to a human gene. We then normalized the matrix to have each column sum to one, effectively creating a count-preserving probabilistic map from d mouse to D human genes M ∈R^D×d^. Mapping from mouse to human genes is then performed with matrix multiplication: U_human_= MU_mouse_. Note that while the mouse gene expression matrix U_mouse_ contains only integers (U_mouse_∈Z^d×N^), the many-to-many mapping means that the mapped human gene expression matrix U_human_ may contain non-integers (U_human_∈R^D×N^). For any human orthologs that were missing in the mouse expression data, we filled in the expression with zeroes. We then log(CP10K+1) normalized the query cells.

#### Preprocessing reference scRNA-seq data

The pancreas reference datasets were each sequenced with a different technology: Fluidigm C1 (n=638 cells), CEL-seq (946 cells), CEL-seq2 (2,238 cells), Smart-seq2 (2,355 cells). We obtained the log(1+CP10K) normalized data from the Harmony publication^17^. The pancreas cells were previously assigned across 9 types within each dataset individually according to cluster-specific expression of marker genes: alpha (*GCG*), beta (*MAFA*), gamma (*PPY*), delta (*SST*), acinar (*PRSS1*), ductal (*KRT19*), endothelial (*CDH5*), stellate (*COL1A2*), and immune (*PTPRC*). We removed 290 cells that were left unassigned as part of ambiguous or outlier clusters during within-dataset annotation, leaving 5,887 reference cells.

We benchmarked three reference mapping methods as follows:

#### Symphony mapping onto a Harmony reference

We calculated the top 1,000 variable genes within each of the four reference dataset separately using VST then pooled them (total 2,236 variable genes) for PCA. For reference integration, we ran Harmony on the top 20 PCs, harmonizing over ‘donor’ (*θ*= 2) and ‘technology’ (*θ*= 4), with *τ*= 5. For Symphony mapping, we specified query ‘donor’, ‘species’, and ‘technology’ covariates.

As a comparison with *de novo* integration, we ran Harmony integration on all 5 datasets together. We pooled the top 1,000 variable genes within each dataset (total 2,650 genes), calculated the top 20 PCs, and harmonized over ‘species’ (*θ*= 2), ‘donor’ (*θ*= 2), and ‘technology’ (*θ*= 2).

#### Seurat v4 mapping onto a Seurat reference

We ran Seurat version 4 (beta)^30^ (Seurat_3.9.9.9024) and followed the steps from the author’s tutorial (https://satijalab.org/seurat/v3.2/integration.html) to integrate the reference datasets given that the *FindIntegrationAnchors* and *IntegrateData* functions for de *novo* integration are equivalent between Seurat v3 and v4 to our understanding. We used the same 2,236 variable genes as above and 20 PCs. We followed the tutorial (https://satijalab.org/seurat/v4.0/reference_mapping.html) to map each donor dataset from the query individually. We used the *FindTransferAnchors* function with reduction = ‘pcaproject’ and *MapQuery* function with reference.reduction = ‘pca’ (as the documentation recommends for unimodal analysis).

As a comparison with *de novo* integration, we ran Seurat v3/4 integration (*FindIntegrationAnchors* and *IntegrateData*) on all 5 datasets (integrating over plate-based technologies and Baron donors as batches) with the same 2,650 variable genes as above.

#### scArches mapping onto a trVAE reference

We ran scArches^28^ version 0.3 with trVAE^33^ using default parameters provided in the authors’ notebooks (https://github.com/theislab/scarches/tree/master/notebooks). For the pancreas analysis, we only had access to normalized expression data and therefore ran scArches with trVAE using the mse reconstruction loss function. We included query batch information in the condition_key parameter.

As a comparison with *de novo* integration, we ran trVAE on all 5 datasets with default parameters, specifying batch as ‘dataset’ for the 4 plate-based datasets and ‘donor’ for the Baron et al. dataset.

#### Evaluation metrics

We used the resulting joint (reference and query) cell embedding to predict query cell types from reference cells using a 5-NN classifier and calculated cell type prediction F1 scores, as described above. Note that for the cell type prediction and cell type F1 score calculation, we excluded query Schwann cells from the accuracy metrics because that cell type is not present in the reference.

To assess degree of mixing, we calculated ref_query LISI and query donor LISI on query cell neighborhoods using the *compute_lisi* function as above. ref_query LISI measures how well the reference and query datasets are mixed (max ref_query LISI = 2), whereas query donor LISI measures how well the individual donors within the query dataset are mixed (max = 6).

We measured mapping runtime and corresponding *de novo* integration runtime for each method as elapsed time starting from gene expression. Symphony and Seurat were run in interactive Jupyter notebooks on a Linux server (Intel Xeon E5-2690 v.3 processors), whereas scArches/trVAE was run on GPUs (graphics card GP100GL [Tesla P100 PCIe 16GB]) to speed up runtime.

### 2.3 Fetal liver hematopoiesis trajectory inference example

We obtained post-filtered, post-doublet removal data directly from the authors^46^ along with author-defined cell type annotations for 113,063 cells sequenced with 10x 3’ end bias and a separate 25,367 cells sequenced with 10x 5’ end bias. For building the harmonized reference from all 3’ cells, we followed the same variable gene selection procedures as the original authors, using the Seurat variance/mean ratio (VMR) method with parameters min_expr = .0125, max_expr = 3, and min_dispersion = 0.625 (resulting in 1,917 variable genes). For each of 14 held-out donor experiments within the 3’ dataset, we integrated the reference with Harmony on 13 donors (*θ*= 3). During Symphony mapping, we specified query ‘donor’ covariate. For mapping 5’ cells against a 3’ reference, we removed two donors (F2 and F5, n=3,953) from the 5’ query based on low library complexity (**Fig. S5b**), leaving n=21,414 cells from 5 donors. We integrated the reference (all 14 donors sequenced with 3’ end bias) with Harmony over ‘donor’ (*θ*= 3). During Symphony mapping, we specified both ‘donor’ and ‘technology’ as covariates. We predicted query cell types by transferring reference cell type annotations using the *knn* function in the ‘class’ R package (k=30). We visualized the aggregated confusion matrix across all 14 held-out donor experiments as well as the confusion matrix for the single 5’-to-3’ experiment using ComplexHeatmap R package^57^.

For the trajectory inference analysis, we obtained trajectory coordinates from the force directed graph (FDG) embedding of all 3’-sequenced cells from the original authors^46^, forming a reference trajectory. We restricted the trajectory to immune cell types only (excluding hepatocytes, fibroblasts, and endothelial). We then mapped a subset of the query cells belonging to the MEM lineage (MEMPs, megakaryocytes, mast cells, early-late erythroid; n=5,141) to the reference-defined trajectory by averaging the FDG coordinates of the 10 reference immune cell neighbors in the Symphony embedding.

### 2.4 Memory T cell surface protein inference example

We used a memory T cell CITE-seq dataset collected from a tuberculosis disease progression cohort of 259 individuals of admixed Peruvian ancestry^40^. The dataset includes expression of the whole transcriptome (33,538 genes) and 30 surface protein markers from 500,089 memory T cells isolated from PBMCs. Including technical replicates, 271 samples were processed across 46 batches.

To assess protein prediction accuracy using Symphony embeddings, we randomly selected 217 samples (411,004 cells), normalized the expression of each gene (log2(CP10K)) and built a Symphony reference based on mRNA expression, correcting for donor and batch. The held-out 54 samples comprised the query that we mapped onto the reference. We predicted the expression of each of the 30 surface proteins in each of the query cells by averaging the protein’s expression across the cell’s 50 nearest reference neighbors. Nearest neighbors were defined based on Euclidean distance in the batch-corrected low-dimensional embedding. As a ground truth for each protein in each query cell, we computed a smoothed estimate of the cells’ measured protein expression by averaging the protein’s expression across the cell’s 50 nearest neighbors in the batch-corrected complete PCA embedding of all 259 donors. We did not use the cells’ raw measured protein expression due to dropout. We computed the Pearson correlation coefficient between our predicted expression and the ground truth expression across all cells per donor for each marker.

To assess protein prediction accuracy based on mapping to a joint mRNA and protein-based Symphony reference, we first built an integrated reference by using canonical correlation analysis (CCA) to project cells into a low-dimensional embedding maximizing correlation between mRNA and protein features. We randomly selected 217 samples (395,373 cells) to comprise this reference, and normalized the expression of each gene (log2(CP10K)), selected the top 2,865 most variable genes, and scaled (mean = 0, variance = 1) all mRNA and protein features. We computed 20 canonical variates (CVs) with the *cc* function in the CCA R package^58^ and corrected the mRNA CVs for donor and batch effects with Harmony. Then, we used Symphony to construct a reference based on the batch-corrected CVs, gene loadings on each CV, and mean and standard deviation used to scale each gene prior to CCA. The held-out 54 samples comprised the query that we mapped onto the reference. As described above, we predicted the expression of each of the 30 surface proteins in each of the query cells based on the cell’s 5, 10, or 50 nearest neighbors in the reference, estimated the smoothed ground truth expression of each protein in each query cell (now based on the batch-corrected CCA embedding of all 259 donors) and computed the Pearson correlation coefficient for each marker.

### 2.5 Visualization

For visualizing the embeddings using UMAP^59^ (and included as the default in Symphony), we used the ‘uwot’ R package with the following parameters: n_neighbors=30, learning_rate=0.5, init = ‘laplacian’, metric = ‘cosine’, min_dist=0.1 (except min_dist=0.3 for pancreas and fetal liver examples). For each Symphony reference, we saved the uwot model at the time of UMAP using the *uwot::save_uwot* function and saved the path to the model file as part of the Symphony reference object. Saving the reference UMAP model allows for the fast projection of new query cells into reference UMAP space from the query embedding from Symphony mapping using the function *uwot::transform*.

For the pancreas benchmarking, we computed a *de novo* UMAP embedding on the joint reference and query embedding because a UMAP projection can potentially obscure differences between the projected data and dataset used to construct the UMAP model. For general purposes, we recommend UMAP projection when the reference cell UMAP coordinates are desired to remain stable.

To distinguish the reference plots from query plots, we visually present the reference embedding as a contour density instead of individual cells. The density plots were generated using ggplot2 function *stat_density_2d* with geom = ‘polygon’ and contour_var = ‘ndensity’. We provide custom functions to generate these plots as part of the Symphony package.

### 2.6 Runtime scalability analysis

We downsampled a large memory T cell dataset^40^ to create benchmark reference datasets with 20,000, 50,000, 100,000, 250,000, and 500,000 cells.For each, we built a reference (20 PCs, 100 centroids) integrating over ‘donor’ and mapped three different-sized queries: 1,000, 10,000, and 100,000 cells. To isolate the separate effects of number of query cells and number of query batches on mapping time, we mapped against the 50,000-cell reference: (1) varying the number of query cells (from 1,000 to 10,000 cells) while keeping the number of donors constant and (2) varying the number of query donors (6 to 120 donors) while keeping the number of cells constant (randomly sampling 10,000 cells). We also performed separate experiments varying the number of reference centroids (25 to 400) and number of dimensions (10 to 320 PCs) while keeping all other parameters constant. We ran all jobs on Linux servers allotted 4 cores and 64 GB of memory (Intel Xeon E5-2690 v.3 processors) and used the *system*.*time* R function to measure elapsed time.

## Data availability

Datasets for all analyses were obtained from the links in **Table S1**. All datasets are publicly available except the memory T cell CITE-seq data, which will be available at GEO accession GSE158769.

## Code availability

We provide an efficient implementation of Symphony at https://github.com/immunogenomics/symphony along with documentation, tutorials, and pre-built references. Scripts reproducing figures for all examples will be made available at https://github.com/immunogenomics/symphony_reproducibility.

