## Supplementary Equations for "Efficient and precise single-cell reference atlas mapping with Symphony"

During Symphony reference building, we calculate two reference compression terms, $N_r$ and $C$, which are precomputed in advance to be used later during the correction step of the reference mapping algorithm. This section describes the relevant parts of the linear mixture model framework shared by Harmony and Symphony and derives the reference compression terms.

### Harmony mixture of experts model learned from reference cells

In the Harmony mixture model learned during reference integration, the *k* clusters represent *k* "experts" in a mixture of experts that serve as surrogate variables for cell states within the low-dimensional space. For each reference cluster $k$, we learn a cluster-specific linear model for each PC: that is, the location in PC space for each reference cell *i* ($Z_{r[i]}$) can be modeled as in (1). For each cluster *k*, we estimated $B_{rk} \in \mathbb{R}^{(1+b)\times d}$, representing the parameters of the linear model. The batch-independent intercept terms $B_{rk[0,\cdot]}$ represent the location of cluster centroid *k* in PC space. The remaining batch-dependent terms $B_{rk[1:b,\cdot]}$ represent reference batch effect coefficients for each PC. See Korsunsky et al. (2019) for full details.

$$Z_{r[i]} = \sum_k R_{r[k,i]} \left[B^T_{rk[0,\cdot]} + B^T_{rk[1:b,\cdot]} X_r\right] + \varepsilon \tag{1}$$

After Harmony integration, the batch effects for the reference cells have been removed by subtracting the batch-dependent terms from each cell. In the final integrated embedding, the harmonized PCs for each reference cell are thereby modeled as the weighted summation of only the intercept terms for the clusters over which the cell is assigned (captured in $R_r$) as well as a cell-specific residual $\varepsilon$.

$$\hat{Z}_{r[i]} = \sum_k R_{r[k,i]} B^T_{rk[0,\cdot]} + \varepsilon \tag{2}$$

### Symphony models non-harmonized query cells with harmonized reference cells

The goal of reference mapping is to add the query cells to our model, modeling all cells together in order to estimate and remove the query batch effects. Let $N = m + n$ represent the total number of cells (sum of number of query and reference cells). Let $X^* \in [0,1]^{(1+c)\times N}$ denote a design matrix for reference mapping in which the first *m* columns represent query cells, and the remaining *n* columns represent harmonized reference cells. The star (*) indicates the design matrix has been augmented: the first row ($X_{[0,\cdot]}$) consists entirely of 1s, corresponding to the batch-independent intercepts (we model the intercepts for all cells). The remaining *c* rows ($X^*_{[1:c,\cdot]}$) represent the one-hot batch assignment of the cells among the *c* query batches. Note that for the reference cell columns, these values are all 0 since the reference cells do not belong to any *query* batches. We do not include reference batch terms in our design matrix because the reference batch-dependent factors have already been removed during reference integration. Therefore, each harmonized reference cell is modeled only by a weighted average of the centroid locations for the clusters over which it belongs and a cell-specific residual.

Let matrix $R \in \mathbb{R}^{k \times N}$ denote the assignment of query and reference cells (columns) across the reference clusters (rows). Then, the parameters ($B_{qk}$) of the mixture of experts model can be solved for as in (3). The notation $diag(R_k) \in \mathbb{R}^{N \times N}$ denotes the diagonalized $k$th row of $R$. Let $Z \in \mathbb{R}^{d \times N}$ denote the horizontal matrix concatenation of the uncorrected query cells in original PC space ($Z_q$) and corrected reference cells in harmonized space ($\hat{Z}_r$). For each cluster *k*, let matrix $B_{qk} \in \mathbb{R}^{(1+c) \times d}$ represent the query parameters to be estimated. The first row of $B_{qk}$ represents the batch-independent intercept terms, and the remaining *c* rows of $B_{qk}$ represent the query batch-dependent coefficients to be estimated.

$$B_{qk} \approx (X^* \, diag(R_k) \, X^{*T} + \lambda I)^{-1} X^* \, diag(R_k) \, Z^T \qquad (3)$$

**Derivation of cached reference-dependent terms**

Instead of directly solving (3) above, we rewrite $diag(R_k)$, $X^*$, and $Z$ by separating out the reference and query-dependent components of each matrix. This allows us to determine which components of the calculation can be precomputed during reference building to reduce computational steps during reference mapping. Assuming the query cells are placed in the first $m$ columns of $R$ and the reference cells are placed in the last $n$ columns of $R$, then $R$ is the horizontal concatenation of $R_q$ and $R_r$. Let vector $R_{q[k,\cdot]}$ of size *m* denote the *k*th row of $R_q$, and let $R_q^{(k)}$ denote the diagonalized square matrix (of dimensions $m \times m$) of $R_{q[k,\cdot]}$. Let vector $R_{r[k,\cdot]}$ of size *n* denote the *k*th row of $R_r$, and let $R_r^{(k)}$ denote the diagonalized square matrix (of dimensions $n \times n$) of $R_{r[k,\cdot]}$. Then, $diag(R_k)$ can be rewritten as $R_q^{(k)} \oplus R_r^{(k)}$, the direct sum of the diagonal matrices for the query and reference cells.

$$diag(R_k) = R_q^{(k)} \oplus R_r^{(k)} = \begin{bmatrix} R_{[k,1]} & \cdots & 0 \\ \vdots & \ddots & \vdots \\ 0 & \cdots & R_{[k,m+n]} \end{bmatrix}$$

$$R_q^{(k)} = \begin{bmatrix} R_{q[k,1]} & 0 & 0 \\ 0 & \ddots & 0 \\ 0 & 0 & R_{q[k,m]} \end{bmatrix} \qquad R_r^{(k)} = \begin{bmatrix} R_{r[k,1]} & 0 & 0 \\ 0 & \ddots & 0 \\ 0 & 0 & R_{q[k,n]} \end{bmatrix}$$

Similarly, we can partition the full design matrix $X^*$ into the left $m$ columns and right $n$ columns that represent the query and reference components: $X_q^* \in \{0,1\}^{(1+c)\times m}$ and $X_r' \in \{0,1\}^{(1+c)\times n}$. The horizontal concatenation of $X_q^*$ and $X_r'^*$ yields $X^*$. Note that $X_r'$ is not the original reference design matrix across reference batches ($X_r$), but rather assignment of reference cells (columns) to *query* batches (rows). Since reference cells do not belong to any query batches, $X_r'$ is a zero matrix, and $X_r'^*$ is the same zero matrix augmented with a row of 1s. In a simple example where there are two batches in the query ($c = 2$), the design matrices take the form:

$$X^* = \begin{bmatrix} 1 \cdots 1 \\ X_q \quad X_r' \end{bmatrix} = \begin{bmatrix} 1 & \cdots & 1\,1 & \cdots & 1 \\ 1 & \cdots & 0\,0 & \cdots & 0 \\ 0 & \cdots & 1\,0 & \cdots & 0 \end{bmatrix}$$

$$X_q^* = \begin{bmatrix} 1 & \cdots & 1 \\ 1 & \cdots & 0 \\ 0 & \cdots & 1 \end{bmatrix} \qquad X_r'^* = \begin{bmatrix} 1 & \cdots & 1 \\ 0 & \cdots & 0 \\ 0 & \cdots & 0 \end{bmatrix}$$

Similarly, we can partition the embedding $Z$ into the left $m$ columns and right $n$ columns that represent the query and reference components: $Z_q \in \mathbb{R}^{d\times m}$ and $\hat{Z}_r \in \mathbb{R}^{d\times n}$, respectively. The horizontal concatenation of $Z_q$ and $\hat{Z}_r$ yields $Z$.

$$Z_q = \begin{bmatrix} Z_{q[1,1]} & \cdots & Z_{q[1,m]} \\ \vdots & \ddots & \vdots \\ Z_{q[d,1]} & \cdots & Z_{q[d,m]} \end{bmatrix} \qquad \hat{Z}_r = \begin{bmatrix} \hat{Z}_{r[1,1]} & \cdots & \hat{Z}_{r[1,n]} \\ \vdots & \ddots & \vdots \\ \hat{Z}_{r[d,1]} & \cdots & \hat{Z}_{r[d,n]} \end{bmatrix}$$

We can then explicitly substitute the query-dependent and reference-dependent components of $diag(R_k)$, $X^*$, and $Z$ to rewrite equation (3) as follows.

$$B_{qk} = \left( [X_q^* \quad X_r'^*] \begin{bmatrix} R_q^{(k)} & 0 \\ 0 & R_r^{(k)} \end{bmatrix} \begin{bmatrix} X_q^T \\ X_r^T \end{bmatrix} + \lambda I \right)^{-1} [X_q^* \quad X_r'^*] \begin{bmatrix} R_q^{(k)} & 0 \\ 0 & R_r^{(k)} \end{bmatrix} \begin{bmatrix} Z_q^T \\ \hat{Z}_r^T \end{bmatrix}$$

$$B_{qk} = \left( \left[ X_q^* R_q^{(k)} \quad X_r'^* R_r^{(k)} \right] \begin{bmatrix} X_q^T \\ X_r^T \end{bmatrix} + \lambda I \right)^{-1} \left[ X_q^* R_q^{(k)} \quad X_r'^* R_r^{(k)} \right] \begin{bmatrix} Z_q^T \\ \hat{Z}_r^T \end{bmatrix}$$

$$B_{qk} = \left( \left[ X_q^* R_q^{(k)} X_q^{*T} + \boldsymbol{X_r'^* R_r^{(k)} X_r'^{*T}} \right] + \lambda I \right)^{-1} \left[ X_q^* R_q^{(k)} Z_q^T + \boldsymbol{X_r'^* R_r^{(k)} \hat{Z}_r^T} \right] \tag{4}$$

In (4), the bolded terms designate terms that depend only on reference cells and can therefore be precomputed ahead of time during the reference building process and subsequently cached for later use during query mapping. The first of these terms, $X_r'^{*} R_r^{(k)} X_r'^{*T}$ of dimensions $(1+c) \times (1+c)$, can be further simplified as follows.

$$\boldsymbol{X_r'^{*} R_r^{(k)} X_r'^{*T}} = \begin{bmatrix} 1 & \cdots & 1 \\ 0 & \cdots & 0 \\ 0 & \cdots & 0 \end{bmatrix} R_r^{(k)} \begin{bmatrix} 1 & 0 & 0 \\ \vdots & \vdots & \vdots \\ 1 & 0 & 0 \end{bmatrix}$$

$$= \begin{bmatrix} R_{r[k,1]} & \cdots & R_{r[k,n]} \\ 0 & \cdots & 0 \\ 0 & \cdots & 0 \end{bmatrix} \begin{bmatrix} 1 & 0 & 0 \\ \vdots & \vdots & \vdots \\ 1 & 0 & 0 \end{bmatrix} = \begin{bmatrix} \sum_{i=1}^{n} R_{r[k,i]} & 0 & 0 \\ 0 & 0 & 0 \\ 0 & 0 & 0 \end{bmatrix} = \begin{bmatrix} N_k & 0 & 0 \\ 0 & 0 & 0 \\ 0 & 0 & 0 \end{bmatrix}$$

Intuitively, $N_k \in \mathbb{R}$ is the number of cells (can be a non-integer number since the cells are soft assigned) belonging to cluster $k$. Therefore, to capture this term for all *k* clusters, we need only save $N_r \in \mathbb{R}^{k \times 1}$, a vector containing the size of each of the $k$ clusters in terms of the number of cells contained within them. Similarly, the second of the reference-dependent terms, $X_r'^{*} R_r^{(k)} \hat{Z}_r^T$, can also be further simplified as follows.

$$\boldsymbol{X_r'^{*} R_r^{(k)} \hat{Z}_r^T} = \begin{bmatrix} 1 & \cdots & 1 \\ 0 & \cdots & 0 \\ 0 & \cdots & 0 \end{bmatrix} R_r^{(k)} \begin{bmatrix} \hat{Z}_{r[1,1]} & \cdots & \hat{Z}_{r[d,1]} \\ \vdots & \ddots & \vdots \\ \hat{Z}_{r[1,n]} & \cdots & \hat{Z}_{r[d,n]} \end{bmatrix}$$

$$= \begin{bmatrix} R_{r[k,1]} & \cdots & R_{r[k,n]} \\ 0 & \cdots & 0 \\ 0 & \cdots & 0 \end{bmatrix} \begin{bmatrix} \hat{Z}_{r[1,1]} & \cdots & \hat{Z}_{r[d,1]} \\ \vdots & \ddots & \vdots \\ \hat{Z}_{r[1,n]} & \cdots & \hat{Z}_{r[d,n]} \end{bmatrix}$$

$$= \begin{bmatrix} R_{r[k,\cdot]} \cdot \hat{Z}_{r[\cdot,1]}^T & \cdots & R_{r[k,\cdot]} \cdot \hat{Z}_{r[\cdot,d]}^T \\ 0 & \cdots & 0 \\ 0 & \cdots & 0 \end{bmatrix}$$

Therefore, to capture this term for all *k* clusters, we need only save $C \in \mathbb{R}^{k \times d}$, a matrix containing $k$ rows, where each row consists of the vector $\left[R_{r[k,\cdot]} \cdot \hat{Z}_{r[\cdot,1]}^T \quad \cdots \quad R_{r[k,\cdot]} \cdot \hat{Z}_{r[\cdot,d]}^T\right]$ for the corresponding cluster. We can directly calculate $C = R_r \hat{Z}_r^T$.
