## Supplementary Figures 1-8 for "Efficient and precise single-cell reference atlas mapping with Symphony"

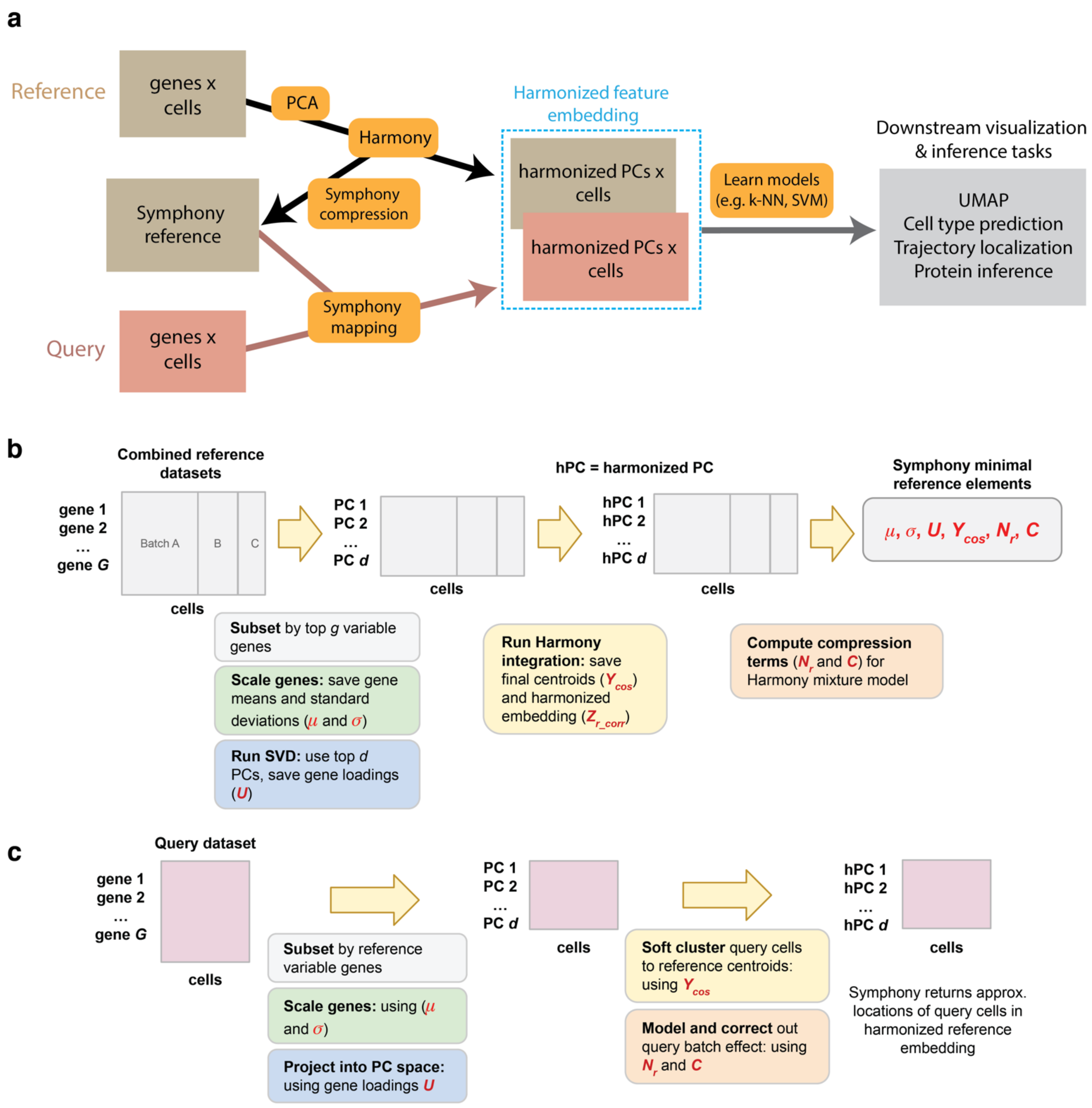

**Supplementary Figure 1. Overview of reference mapping pipeline and Symphony data structures. (a)** The overall analysis pipeline comprises various functions (orange boxes) that each perform a transformation on the data. Symphony mapping takes in a query gene expression matrix and a Symphony reference built from integrated reference datasets, and outputs the query cell locations in the harmonized feature embedding. Models trained on the reference feature embedding (e.g. cell type classifier) can transfer annotations to the query for various downstream tasks. **(b)** Steps of reference building algorithm. Reference datasets spanning multiple batches are aggregated into a single expression matrix on which PCA and Harmony integration is performed. The output of reference compression is the Symphony minimal reference elements, consisting of data structures $\mu, \sigma$, $U$, $Y_{cos}$, $N_r$, and $C$ (red symbols). $Z_{r\_corr}$ (the harmonized reference embedding) is not used for the mapping calculation but is saved for downstream annotation transfer. **(c)** Steps of query mapping algorithm, indicating where each reference element is used. Query cells are projected into reference PCA space, clustered to reference centroids, and corrected to harmonized space by removing query batch effects.

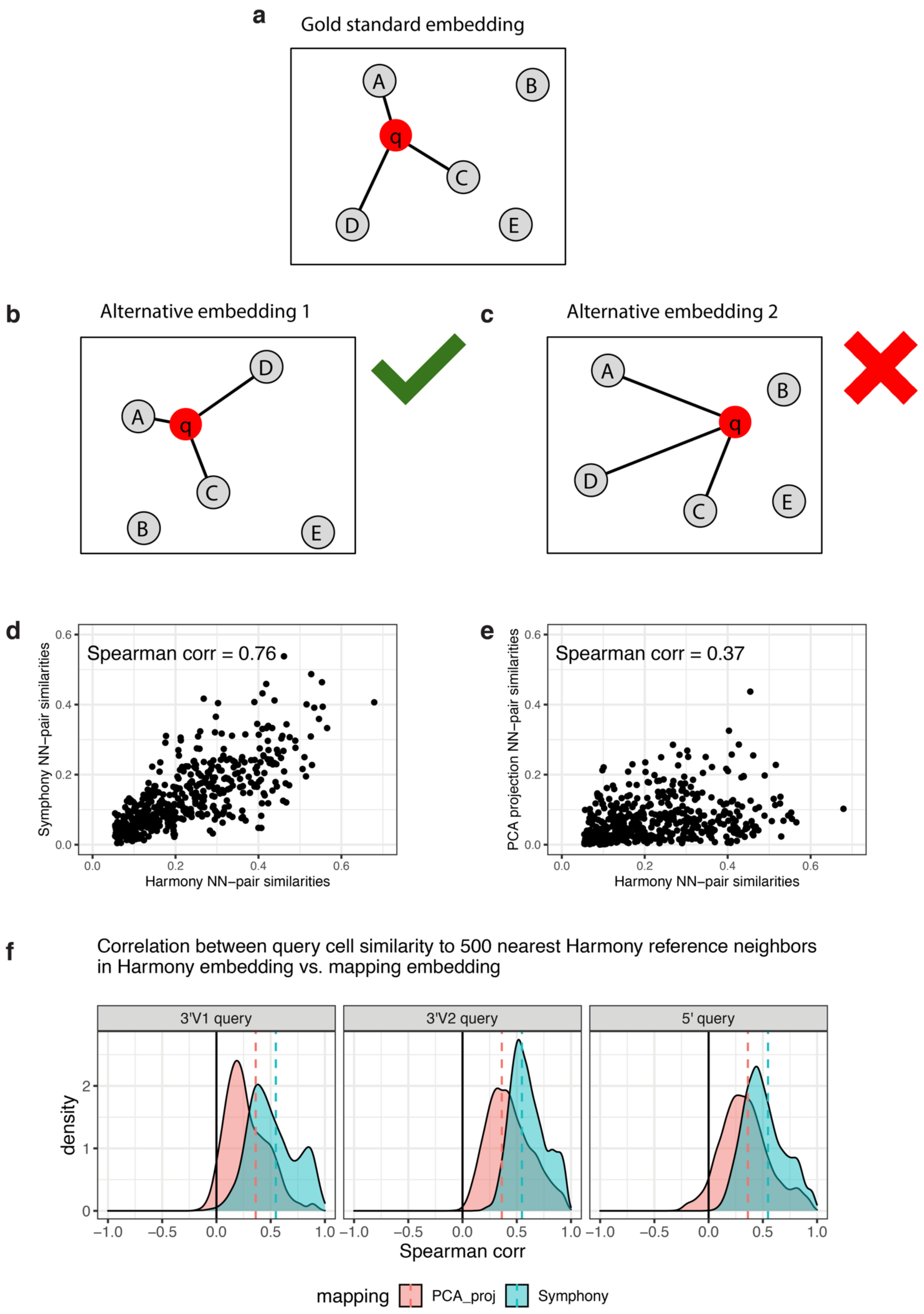
a
Gold standard embedding
A
B
q
C
D
E
b
Alternative embedding 1
D
A
q
C
B
E
c
Alternative embedding 2
A
B
q
D
C
E
d
Spearman corr = 0.76
Symphony NN−pair similarities
Harmony NN−pair similarities
0.0
0.2
0.4
0.6
e
Spearman corr = 0.37
PCA projection NN−pair similarities
Harmony NN−pair similarities
0.0
0.2
0.4
0.6
f
Correlation between query cell similarity to 500 nearest Harmony reference neighbors
in Harmony embedding vs. mapping embedding
3'V1 query
3'V2 query
5' query
density
0
1
2
−1.0
−0.5
0.0
0.5
1.0
Spearman corr
mapping
PCA_proj
Symphony

**Supplementary Figure 2. Nearest neighbor correlation (k-NN-corr) metric.** The k-NN-correlation metric assesses how well an alternative embedding recapitulates the structure of a gold standard embedding. k-NN-corr is asymmetric in that it matters which of the two embeddings is selected as the gold standard. Consider a gold standard embedding **(a)** and two alternative embeddings **(b)** and **(c)**, representing a good mapping and a bad mapping, respectively. For a given query cell *q* (red), we identify its top *k* nearest reference (gray) neighbors in the gold standard embedding (*k* = 3 depicted) and calculate the similarity between the query cell and each neighbor. The similarities between the same query-reference neighbor pairs are then calculated in the alternate embedding. k-NN-corr is the Spearman correlation between the similarities in the gold standard vs. alternative embedding, ranging from -1 to +1. Example k-NN-corr for one query cell and *k* = 500 for the **(d)** Symphony embedding and **(e)** PCA projection embedding. **(f)** k-NN-corr distribution across query cells for k=500 and a gold standard Harmony embedding, for either the Symphony embeddings (blue) or a simple PCA projection with no correction step (red), faceted by query dataset.

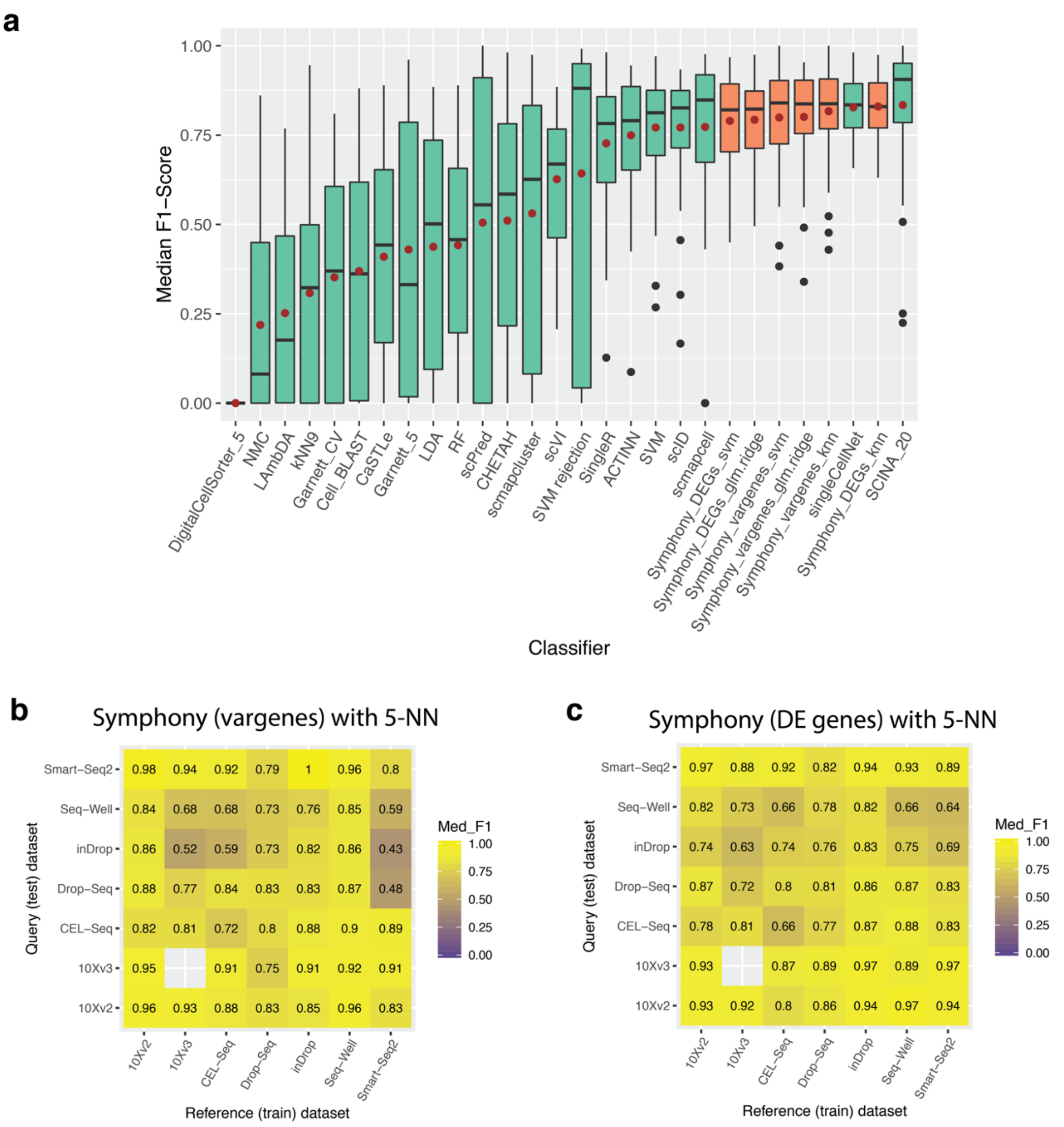


**Supplementary Figure 3. Symphony performance against automatic cell type classifiers.** Following the cross-technology PBMC benchmarking experiment from Abdelaal et al. (2019), we ran a total of 48 train-test experiments per Symphony-based classifier. Two different versions of the Symphony feature embeddings were generated depending on variable gene selection method: top 2000 variable genes (vargenes) or top 20 differentially genes (DEGs) expressed per cell type. Symphony embeddings were used to train 3 downstream classifiers: k-NN (k=5), SVM with radial kernel, and multinomial logistic regression (glmnet) with ridge. **(a)** Symphony (orange) median cell-type F1 score across 48 train-test experiments compared to supervised methods (green), demonstrating noninferiority to the top supervised methods and stable performance regardless of downstream classification method. Red dot indicates mean of median F1 scores across 48 experiments (used for ordering the methods along the x-axis). **(b, c)** Median cell type F1 score across 48 experiments for the 5-NN classifier with variable gene selection **(b)** and DEG selection **(c)**. Non-diagonal values represent train on one technology, test on another (42 experiments, all with donor 1). Values along the diagonal indicate train on donor 1, test on donor 2 of the same technology (6 experiments; missing square because donor 2 not sequenced with 10x v3).

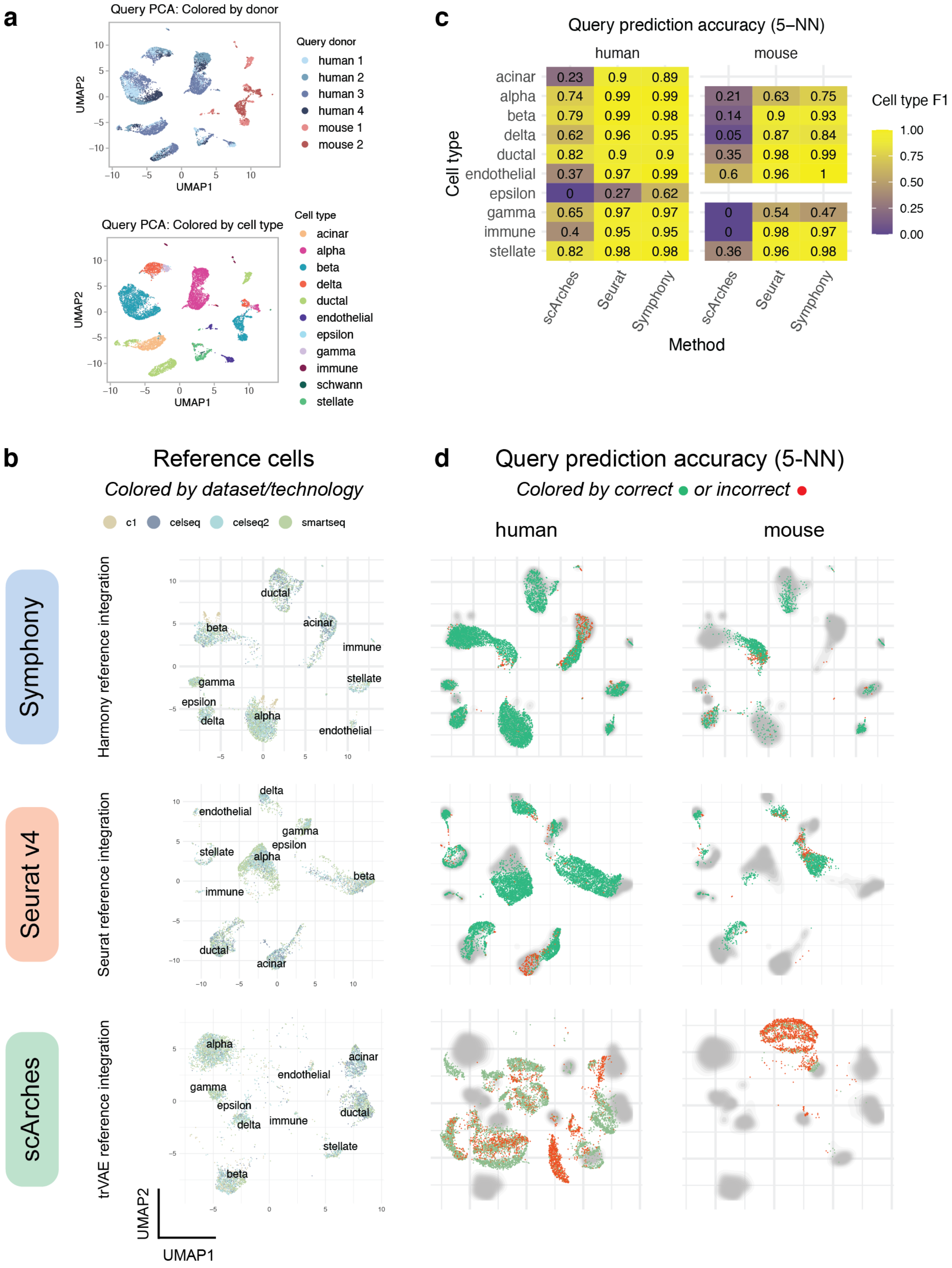
a
Query PCA: Colored by donor
Query donor
human 1
human 2
human 3
human 4
mouse 1
mouse 2
UMAP2
UMAP1
Query PCA: Colored by cell type
Cell type
acinar
alpha
beta
delta
ductal
endothelial
epsilon
gamma
immune
schwann
stellate
c
Query prediction accuracy (5–NN)
human
mouse
Cell type
acinar 0.23 0.9 0.89
alpha 0.74 0.99 0.99 0.21 0.63 0.75
beta 0.79 0.99 0.98 0.14 0.9 0.93
delta 0.62 0.96 0.95 0.05 0.87 0.84
ductal 0.82 0.9 0.9 0.35 0.98 0.99
endothelial 0.37 0.97 0.99 0.6 0.96 1
epsilon 0 0.27 0.62
gamma 0.65 0.97 0.97 0 0.54 0.47
immune 0.4 0.95 0.95 0 0.98 0.97
stellate 0.82 0.98 0.98 0.36 0.96 0.98
scArches
Seurat
Symphony
Method
Cell type F1
1.00
0.75
0.50
0.25
0.00
b
Reference cells
Colored by dataset/technology
c1
celseq
celseq2
smartseq
Symphony
Harmony reference integration
Seurat v4
Seurat reference integration
scArches
trVAE reference integration
UMAP2
UMAP1
d
Query prediction accuracy (5-NN)
Colored by correct or incorrect
human
mouse

**Supplementary Figure 4. Comparison of Symphony to alternative reference mapping methods on a cross-species pancreatic islet cell benchmark. (a)** Standard PCA pipeline applied to the Baron et al. query dataset exhibits strong species and donor effects, demonstrating the need for within-query integration. We benchmarked Symphony mapping (on a Harmony-integrated reference), Seurat v4 mapping (on a Seurat anchor-based-integrated reference), and scArches mapping (on a trVAE-integrated reference). For each approach, we built an integrated reference **(b)**, mapped the query, then predicted query cell types using a 5-NN classifier to transfer annotations using the respective reference embedding. **(c)** Query cell prediction accuracy by species for each method as measured by cell type F1 score, with author-defined ground truth labels. Mouse samples did not have acinar or epsilon cells. The resulting joint cell embedding for each tool was visualized by UMAP **(b, d)**: **(b)** Reference cells colored by dataset/technology. **(d)** Query cells colored by correct (green) or incorrect (red) cell type prediction.

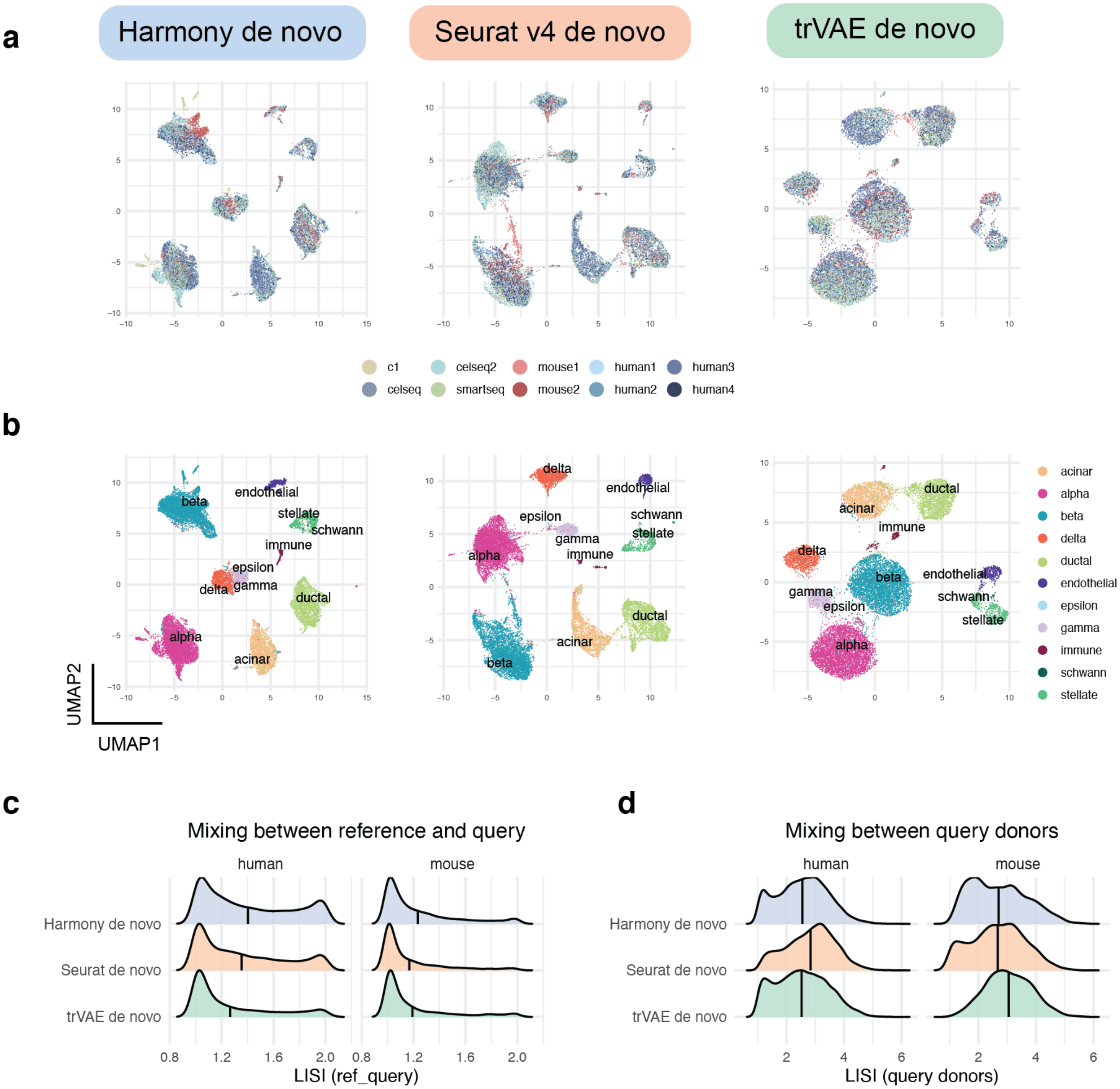


**Supplementary Figure 5. Comparison of *de novo* integration methods for harmonizing all five pancreatic islet cell datasets.** As a comparison to reference mapping (Fig 3), we integrated all five pancreatic islet cell technologies (n=16,342 cells) using 3 *de novo* integration methods: Harmony, Seurat anchor-based integration, and trVAE. UMAP visualizations for the integrated embedding colored by batch **(a)** and cell types **(b)** for each method. Cell types for reference datasets (c1, celseq, celseq2, smartseq) were defined within each dataset separately based on marker genes. Query cell types were defined by Baron et al. Degree of mixing between reference and query datasets **(c)** and mixing between query donors **(d)** was measured with LISI metric on query cell neighborhoods for each method, demonstrating equivalent mixing among *de novo* integration methods (compare to Fig 3d-e).

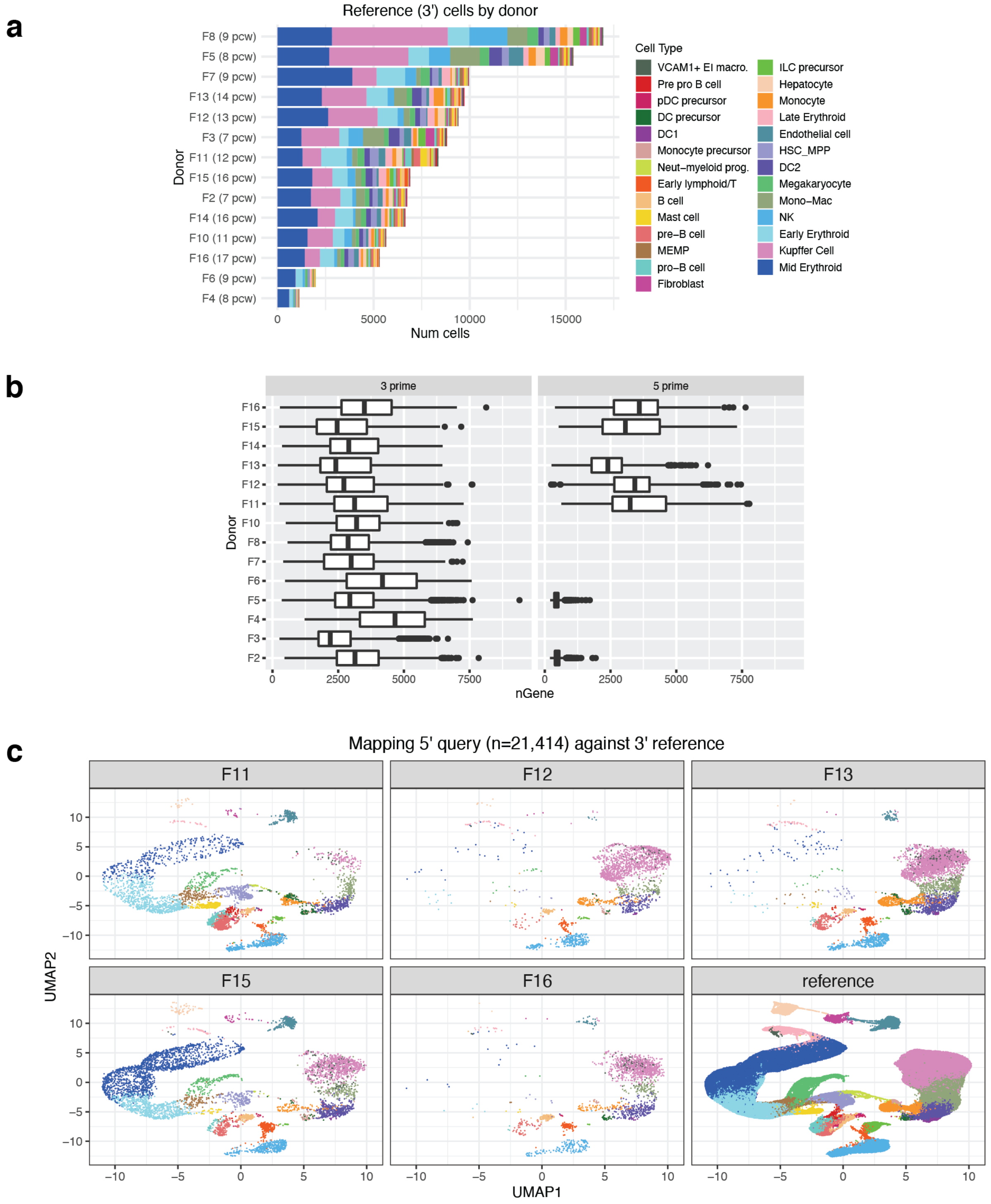


**Supplementary Figure 6. Mapping to a fetal liver hematopoiesis trajectory. (a)** Size and cell type composition of each donor sample in the 10x 3' dataset across 27 author-defined cell types from Popescu et al. (2019). pcw = post-conception weeks. **(b)** Library complexity for each sample in 10x 3' and 10x 5' datasets, showing low complexity for donor F2 and F5 5'-sequenced samples (removed from further analysis). **(c)** UMAP projections of query cells into reference UMAP space after Symphony mapping, faceted by query donor, colored by cell type. Reference UMAP embedding in bottom-right.

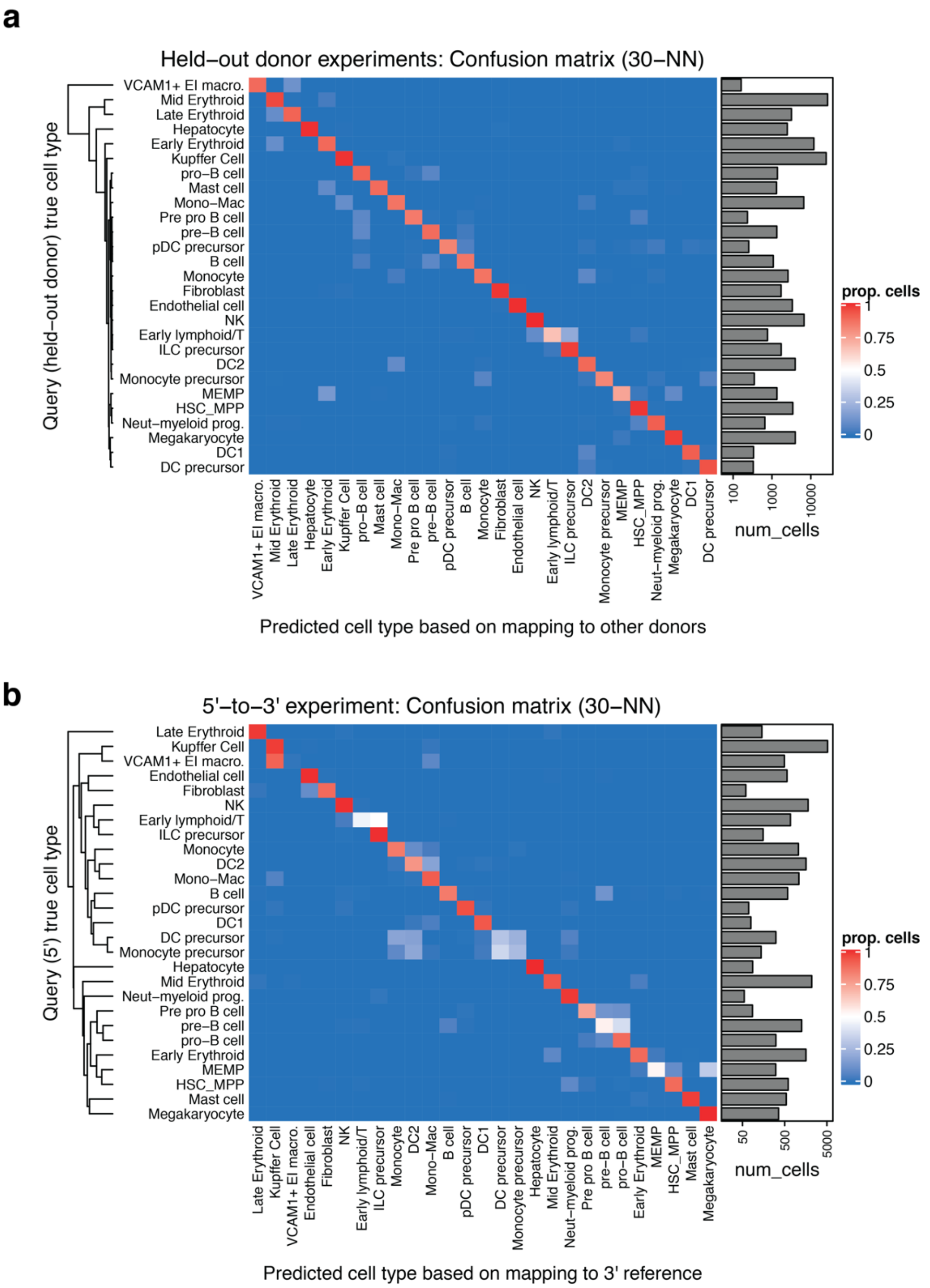


**Supplementary Figure 7. Fetal liver hematopoiesis cell type classification confusion matrices.** We performed two versions of the reference mapping experiments to assess cell type classification accuracy across 27 fine-grained cell types: (1) using exclusively 10x 3' data, we mapped one held-out donor against a reference constructed from the remaining 13 donors (total 14 mapping experiments), (2) mapping all 10x 5' cells against all 10x 3' cells. Cell type confusion matrices are shown for a 30-NN cell type classifier **(a)** aggregated across the 14 held-out donor experiments using exclusively 3' data and **(b)** the 5'-to-3' experiment mapping the full 5' query (n=21,414, n=5 donors) against the full 3' reference (n=113,063 cells, 14 donors), colored by the proportion of the true cell type that was classified correctly. True cell type is defined by the original authors (Popescu et al., 2019). Rows (true query cell types) are sorted by hierarchical clustering on the average gene expression (all genes) for the cell types to order similar types together. Bar graph (right) shows population size for each cell type.

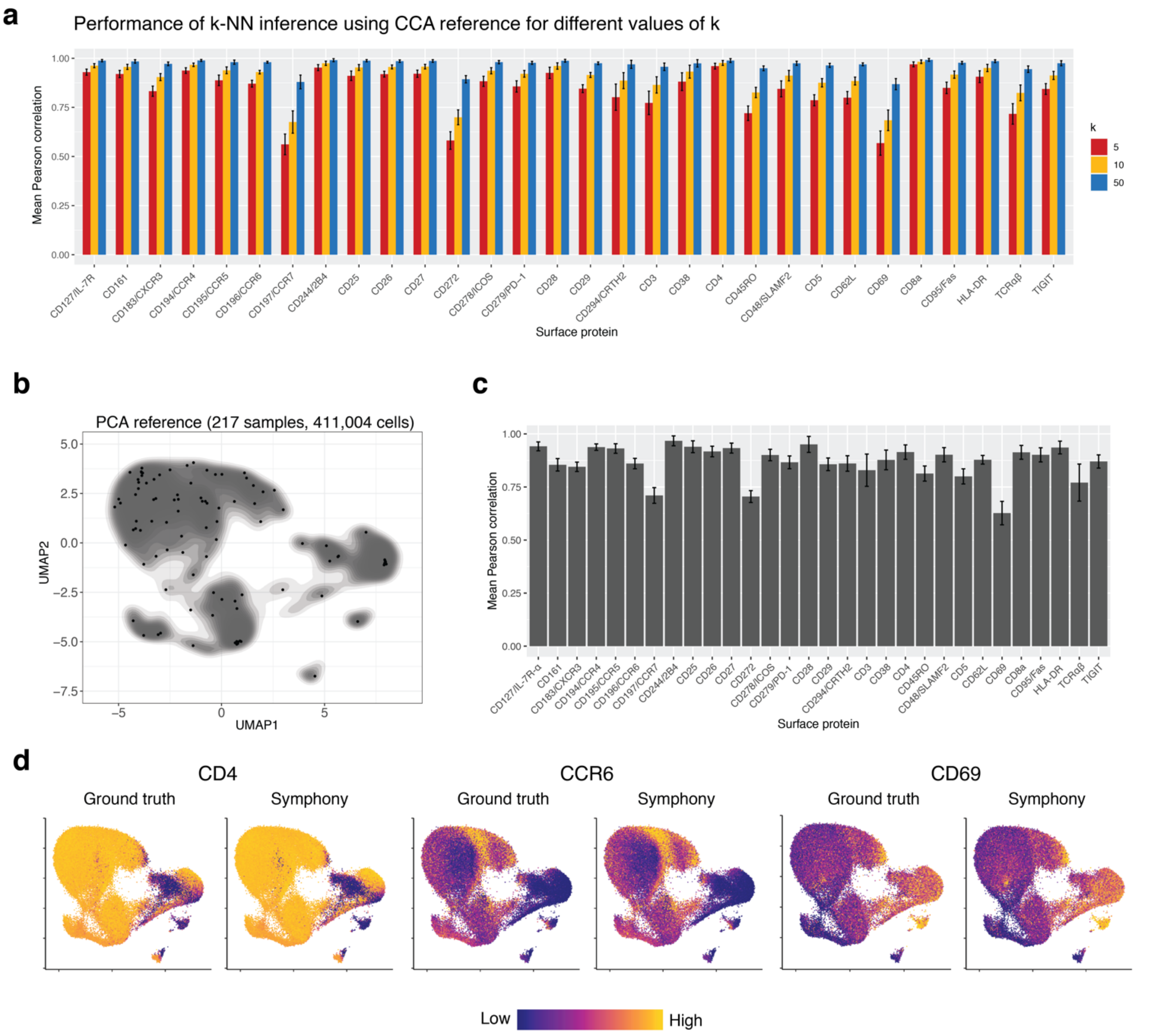


**Supplementary Figure 8. Inferring query surface protein expression in memory T cells. (a)** Mean Pearson correlation for CCA reference between k-NN predicted protein expression and ground truth for different values of *k*. **(b)** Symphony reference built from a standard mRNA PCA embedding (reference protein values were not used to build embedding but treated as annotations only). Contour fill represents density of reference cells. Black points represent soft-cluster centroids in the Symphony mixture model. **(c)** We measured the accuracy of protein expression prediction based on the PCA reference with the Pearson correlation between predicted and ground truth expression for each surface protein across query cells in each donor. Bar height represents the average per-donor correlation for each protein, and error bars represent standard deviation. **(d)** Ground truth and predicted expression of CD4, CCR6, and CD69 based on PCA reference. Ground truth is the 50-NN-smoothed expression measured in the CITE-seq experiment. Colors are scaled independently for each marker from minimum (blue) to maximum (yellow) expression.
